# *MIR3607* regulates cerebral cortex development via activation of Wnt/βCat signaling

**DOI:** 10.1101/729939

**Authors:** Kaviya Chinnappa, Ángel Márquez-Galera, Anna Prieto-Colomina, Yuki Nomura, Adrián Cárdenas, José P. López-Atalaya, Víctor Borrell

## Abstract

The evolutionary expansion of the mammalian cerebral cortex is recapitulated during embryonic development in large mammals, but the underlying genetic mechanisms remain mostly unknown. Previous transcriptomic analyses of the developing ferret cortex identify candidate genes related to the expansion of germinal layers and cortex size. Here we focused on *MIR3607*, a microRNA differentially expressed between germinal layers of the large human and ferret cortex, not expressed in the small mouse cortex. Expression of *MIR3607* in mouse embryos at E14.5 leads to increased progenitor cell proliferation. This is reflected in transcriptomic changes, which also reveal increased Wnt/βCatenin signaling. Expression of *MIR3607* at E12.5, when progenitor cells expand, causes amplification and severe delamination of apical progenitors, leading to rosette formation. This is rescued by co-expressing Adenomatous Polyposis Coli, inhibitor of canonical Wnt signaling. A similar phenotype is produced in human cerebral organoids. Our findings demonstrate that *MIR3607* expands and delaminates apical progenitor cells via activating Wnt/βCatenin, and suggest that a secondary loss of expression in mouse may underlie their reduction in cortex size during recent evolution.

## Introduction

The mammalian cerebral cortex went through a remarkable expansion in size and folding during evolution from its initial ancestor (O’Leary et al., 2013), a process recapitulated during embryonic development (Borrell & Reillo, 2012; Lewitus et al., 2014). At the cellular level, this is the result of the increased pool size of neural stem and progenitor cells, and their proliferative capacity (Dehay & Kennedy, 2007; Fernandez et al., 2016; Rakic, 2009). In some clades like new world monkeys and, particularly, rodents, this process was reversed at some point in evolution: their brains evolved to be smaller and smoother than those of their ancestors (Kelava et al., 2013; Kelava et al., 2012). This seems to have resulted from the secondary loss of developmental features key for brain growth and folding (Borrell & Reillo, 2012; Fernandez et al., 2016). Understanding the cellular and molecular mechanisms of cerebral cortex expansion during mammalian evolution has recently become a topic of widespread interest. Recent studies have identified genes that emerged in the recent human lineage and that promote neural stem cell proliferation and brain growth (Fiddes et al., 2018; Florio et al., 2015; Florio et al., 2017; Suzuki et al., 2018). This is also achieved by the expression of highly conserved genes in new patterns or at new levels (Cardenas et al., 2018). Unfortunately, our understanding of the molecular mechanisms that regulate the different levels of activity of key signaling pathways across mammalian phylogeny and in brain evolution remains limited.

Neural stem and progenitor cells in the developing mouse cerebral cortex are organized in two main germinal zones. The Ventricular Zone (VZ) is the inner (apical) layer of the embryonic cortex and is composed of apical Radial Glia Cells (aRGCs), the primary type of cortical progenitor cell (Malatesta et al., 2000; Noctor et al., 2001). The Subventricular Zone (SVZ) sits basal to the VZ and houses mostly Intermediate Progenitor Cells (IPCs), a secondary type of progenitor cell generated by aRGCs that produces the majority of cortical neurons (Kowalczyk et al., 2009; Miyata et al., 2004; Noctor et al., 2004; Taverna et al., 2014). In species with very large brains like carnivores and primates, the abundance of SVZ progenitor cells increases massively during early embryonic development, and this layer becomes subdivided into Inner and Outer SVZ (ISVZ, OSVZ) (Dehay & Kennedy, 2007; Reillo & Borrell, 2012; Reillo et al., 2011; Smart et al., 2002). In addition to IPCs, ISVZ and OSVZ are very abundant in basal Radial Glia Cells (bRGCs) (Reillo & Borrell, 2012; Reillo et al., 2011). These are progenitor cells similar to aRGCs, generated by them via delamination from the VZ, that contribute critically to increase neurogenesis and to promote cerebral cortex folding in mammals (Fietz et al., 2010; Hansen et al., 2010; LaMonica et al., 2013; Martinez-Martinez et al., 2016; Nonaka-Kinoshita et al., 2013; Reillo et al., 2011; Stahl et al., 2013).

The rates of progenitor cell amplification and neurogenesis are dramatically different between species with large brains (i.e. human, macaque, ferret) and small brains (i.e., mouse) (Betizeau et al., 2013; Dehay & Kennedy, 2007; Reillo & Borrell, 2012). Within large-brained species, differences also exist between cortical regions (Lukaszewicz et al., 2005), and particularly between prospective folds and fissures (Reillo et al., 2011). Primate- and human-specific genes have been identified that promote cortical progenitor cell amplification and neurogenesis, including protein-coding genes (Fiddes et al., 2018; Florio et al., 2018; Suzuki et al., 2018), and microRNAs that target cell cycle proteins (Arcila et al., 2014; Nowakowski et al., 2018). Whereas these genes may contribute to the emergence of primate-specific features (Florio et al., 2017; Nowakowski et al., 2018), they do not explain differences across phylogeny between mammals with big and small brains (i.e., human, ferret versus mouse). On the other hand, transcriptomic analyses of cortical germinal zones have identified differences in expression levels of highly conserved genes: a) between mouse and macaque or human embryos (Arcila et al., 2014; Ayoub et al., 2011; Fietz et al., 2012); b) underlying the formation of bRGCs and the OSVZ (Martinez-Martinez et al., 2016); and c) related to the different expansion of the cerebral cortex between folds and fissures (de Juan Romero et al., 2015). Among these genes, the identification of several miRNAs suggests that they may be key as genetic regulators of cortical expansion across mammals (Dehay et al., 2015; Fernandez et al., 2016; Florio et al., 2017).

Here we focused on *MIR3607-5p* (SNORD138), a small nucleolar RNA originally identified in irradiated human cells (Chaudhry et al., 2013), highly expressed in human lung cancer cells (Lin et al., 2017), and that we previously found expressed as pre-*MIR3607* in the germinal layers of the developing ferret cerebral cortex (de Juan Romero et al., 2015; Martinez-Martinez et al., 2016). We found that mature *MIR3607-5p* is expressed in cortical germinal layers of the developing human and ferret but not mouse embryo, and took advantage of this to study the effect of expressing it on the development of the mouse cerebral cortex. At mid-neurogenesis, *MIR3607* expression initially increased cell cycle re-entry of aRGCs, but one day later it reversed to increasing neurogenesis. Prematurely-born neurons displayed migration defects, either migrating past their termination zone or remaining within the germinal zone to form a subcortical heterotopia. Transcriptomic analyses revealed that *MIR3607* strongly promotes progenitor cell proliferation and stemness, by acting as a key regulator of the canonical Wnt/βCatenin pathway via directly targeting Adenomatous Polyposis Coli (APC), a critical repressor of βCatenin function. Expression of *MIR3607* at early stages of cortical development significantly overactivated βCatenin signaling in mouse embryos, which led to the amplification of aRGCs and severe disruption of the apical adherens junction belt. This was followed by the delamination of aRGCs and formation of proliferative rosettes, which in the postnatal cortex developed into subcortical heterotopia. These phenotypes were rescued by the additional expression of APC, confirming that they resulted from the overactivation of βCatenin signaling by *MIR3607*. Our results identify *MIR3607* as a key regulator of Wnt/βCatenin signaling during cortical development, and its role in the evolutionary expansion of neural stem and progenitor cells and their delamination to basal compartments, underlying the expansion of the mammalian cerebral cortex.

## Results

### Mature *MIR3607* is expressed in the embryonic cerebral cortex of human and ferret but not mouse

Our previous transcriptomic analyses revealed that pre*-MIR3607* is expressed in all germinal layers of the ferret cerebral cortex at mid-to-late stages of cortical development. This includes the critical period for formation of the OSVZ by massive seeding of bRGCs (Martinez-Martinez et al., 2016), and the late period preceding the phenomenal expansion and folding of the cerebral cortex (de Juan Romero et al., 2015). In contrast, pre-*MIR3607* has never been reported to be expressed in the developing mouse cortex, small and smooth. These observations prompted the idea that *MIR3607* might be relevant for the expansion and folding of the mammalian cerebral cortex. Unfortunately, expression of pre-microRNAs does not always parallel miRNA activity, as those must be enzymatically processed and cleaved to the shorter, functionally mature miRNAs. To gain insights into the potential function of *MIR3607* in cortex expansion and folding, we began elucidating the expression level and pattern of the mature *MIR3607*-5p (*MIR3607* from hereon) in the developing human brain. *In situ* hybridization (ISH) stains on the embryonic human cortex at 16 gestational weeks (GW; circa the peak of neurogenesis) revealed the highest expression levels of *MIR3607* in the VZ and ISVZ, and in the cortical plate (Fig. 1A). Expression was also detected in the OSVZ but at lower levels compared to the other germinal zones (Fig. 1A’).

**Figure 1.**
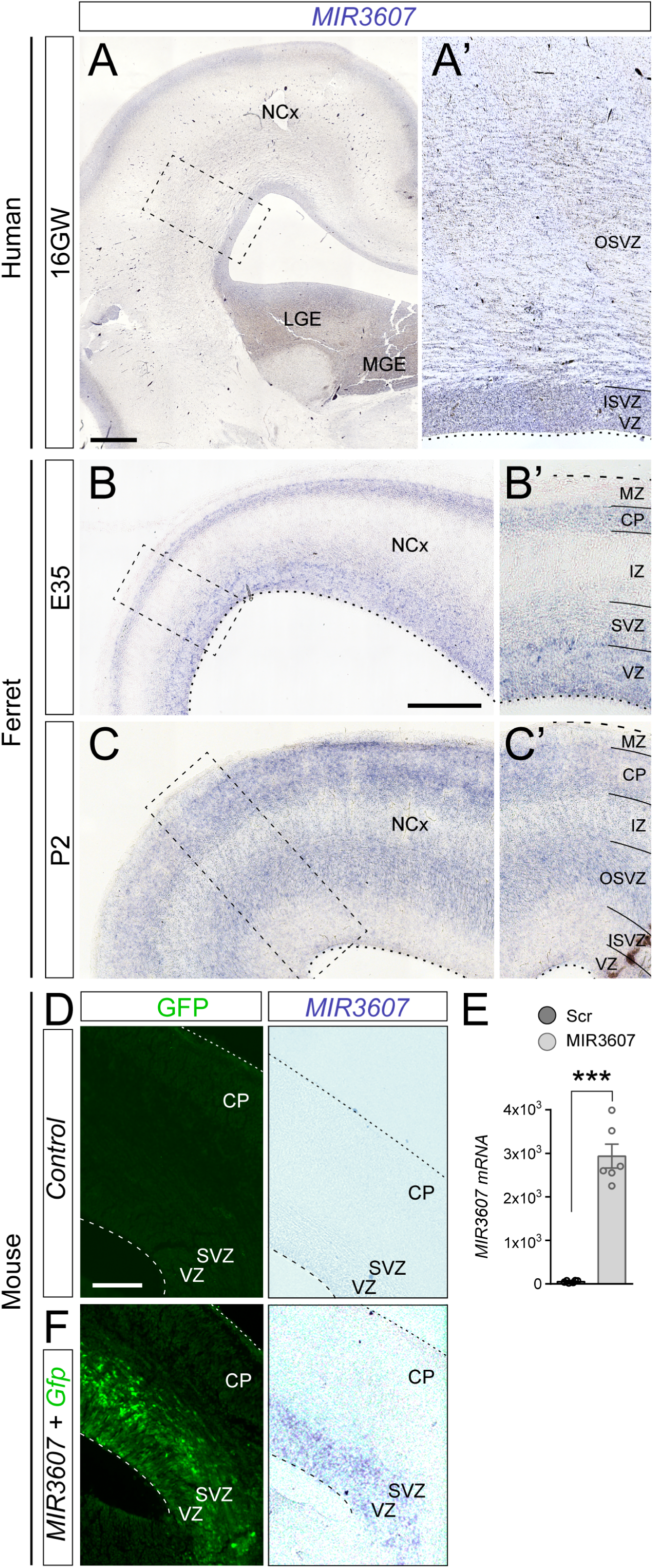
Mature *MIR3607* is expressed in cortical germinal layers of human and ferret but not mouse embryos. (**A,A’**) Coronal section of the human brain at 16 gestational weeks showing the expression pattern of *MIR3607*. Area boxed in (A) is shown in (A’). *MIR3607* expression is high in the three germinal zones: ventricular (VZ), inner and outer subventricular zones (ISVZ and OSVZ, respectively). (**B-C’**) Sagittal sections of the developing ferret cerebral cortex at the indicated ages showing the expression pattern of *MIR3607*. Areas boxed in (B,C) are shown in (B’,C’). At E35, *MIR3607* is expressed at high levels in ventricular zone (VZ) and low levels in subventricular zone (SVZ). At P2, expression is undetectable in VZ and ISVZ, but high in OSVZ. Expression is high in cortical plate (CP) in both human and ferret. (**D,F**) Patterns of GFP and *MIR3607* expression in the cerebral cortex of an E15.5 mouse embryo in a control hemisphere (D), and in the contralateral hemisphere electroporated at E14.5 with *Gfp* plus *MIR3607*-encoding plasmids (F). Note the absence of endogenous expression in (D), and the high expression levels in VZ and SVZ in (F). (**E**) qPCR results of *MIR3607* expression levels (arbitrary units) in HEK293 cells transfected with psil-*Scr* and psil-*MIR3607*. Histograms indicate mean ± SEM; circles in plots indicate values for individual replicas. t-test, \*\*\**p*=9.76×10^-7^. Scale bars: 500µm (A), 200µm (B,C), 100µm (D,F).

Next, we analyzed *MIR3607* expression in the developing ferret cortex. At embryonic day (E) 35, expression was high in VZ, low in SVZ and undetectable in IZ (Fig. 1B,B’). At this stage in ferret, there is no distinction between ISVZ and OSVZ, so this pattern was reminiscent of the difference between VZ and OSVZ in human embryos at 16GW. However, at postnatal day (P) 2, when ISVZ and OSVZ are clearly distinct in ferret, expression was reduced to background levels in VZ and ISVZ, while being high in OSVZ, and intermediate in IZ (Fig. 1C,C’). In addition, and similar to human, *MIR3607* expression was high at both developmental stages in CP, where cortical neurons finish radial migration and begin differentiating their dendritic and axonal arbors. In contrast to human and ferret, our ISH stains revealed a complete absence of *MIR3607* expression in the embryonic mouse cortex (Fig. 1D). Confirmation that *MIR3607* is not expressed in the developing mouse cortex prompted us to take advantage of this circumstance in order to investigate the role of this microRNA in cortex development, and its potential relevance in the limited growth of the mouse cerebral cortex as compared to ferret and human.

MicroRNAs usually have a high degree of conservation across animal species (Gebert & MacRae, 2019). For *MIR3607-5p*, the sequence conservation compared to human ranges between 100% in hominids and 92% in the house mouse (**Supp Fig. S1**). Nucleotide changes mostly occur in the 9^th^ and 6^th^ base of the loop sequence (in 29 and 19 species, respectively, out of 29 species with mismatches), and secondarily in the 4^th^ base of the seed sequence (in 18 out of 29 species; always G to A changes; **Supp Fig. S1**). Outside primates, the mature *MIR3607*-5p sequence is 100% conserved in only 4 of the 26 species compared, which belong to three far-related orders: guinea pig (Rodentia), ferret (Carnivora), wild boar and horse (Artiodactyla). In mouse, the seed sequence is fully conserved, but the first base of the mature *MIR3607*-5p is changed (G to A), and four nucleotides are changed or missing in the loop sequence (**Supp Fig. S1**). The striking similarity of the mature *MIR3607* in human, ferret and mouse is suggestive of a preserved functionality for this miRNA in the mouse brain. Therefore, we reasoned that re-expression of *MIR3607* in the developing brain of mouse embryos may shed light on its role during cortical development. We confirmed expression of mature *MIR3607* from our DNA vector both in a human cell line by RT-qPCR, and in the cerebral cortex of electroporated mouse embryos by ISH (Fig. 1E,F).

### Defects in neurogenesis, neuron migration and axon growth following *MIR3607* expression

The expression patterns of *MIR3607* in the developing cortex of human and ferret supported the notion that it may be involved in multiple steps of cortical development, including VZ progenitor cell proliferation, OSVZ formation and amplification, neurogenesis, radial migration through IZ and neuronal differentiation in CP (Fig. 1). We first studied its potential involvement in neurogenesis and radial migration. Benefitting from the absence of *MIR3607* in mouse, we expressed its mature form alongside a reporter plasmid in apical progenitor cells of the E14.5 mouse cortex by *in utero* electroporation. One day after electroporation (E15.5) the majority of GFP+ cells in control, *Scrambled*-electroporated embryos populated the VZ, with only half as many found in SVZ and very few in IZ (Fig. 2A,B). This distribution of GFP+ cells was similar in embryos expressing *MIR3607*, except for a clear tendency to having fewer cells in IZ and SVZ, while more in VZ (Fig. 2A,B). One day later (E16.5) we observed a 21-35% reduction of GFP+ cells in VZ and SVZ of *MIR3607*-expressing embryos, bound to a 54% increase in IZ, as compared to control embryos (Fig. 2C,D). During normal development, once neurons are born in VZ or SVZ they spend some time within the SVZ undergoing short-distance multipolar migration, and only after that they acquire bipolarity, enter the IZ and undergo fast radial migration to the CP (Martinez-Martinez et al., 2018; Tabata & Nakajima, 2003). Hence, the increase in IZ cells suggested that *MIR3607* expression augmented the relative abundance of radially migrating neurons. We reasoned that this could have two main origins: a greater abundance of neurons due to increased neurogenesis, or a faster migration of newborn neurons into the IZ. Analysis of the rate of cell cycle exit revealed a 29% increase at E16.5 upon *MIR3607* expression from E14.5 (Fig. 2E), demonstrating more self-consuming, neurogenic divisions of progenitor cells at this stage.

**Figure 2.**
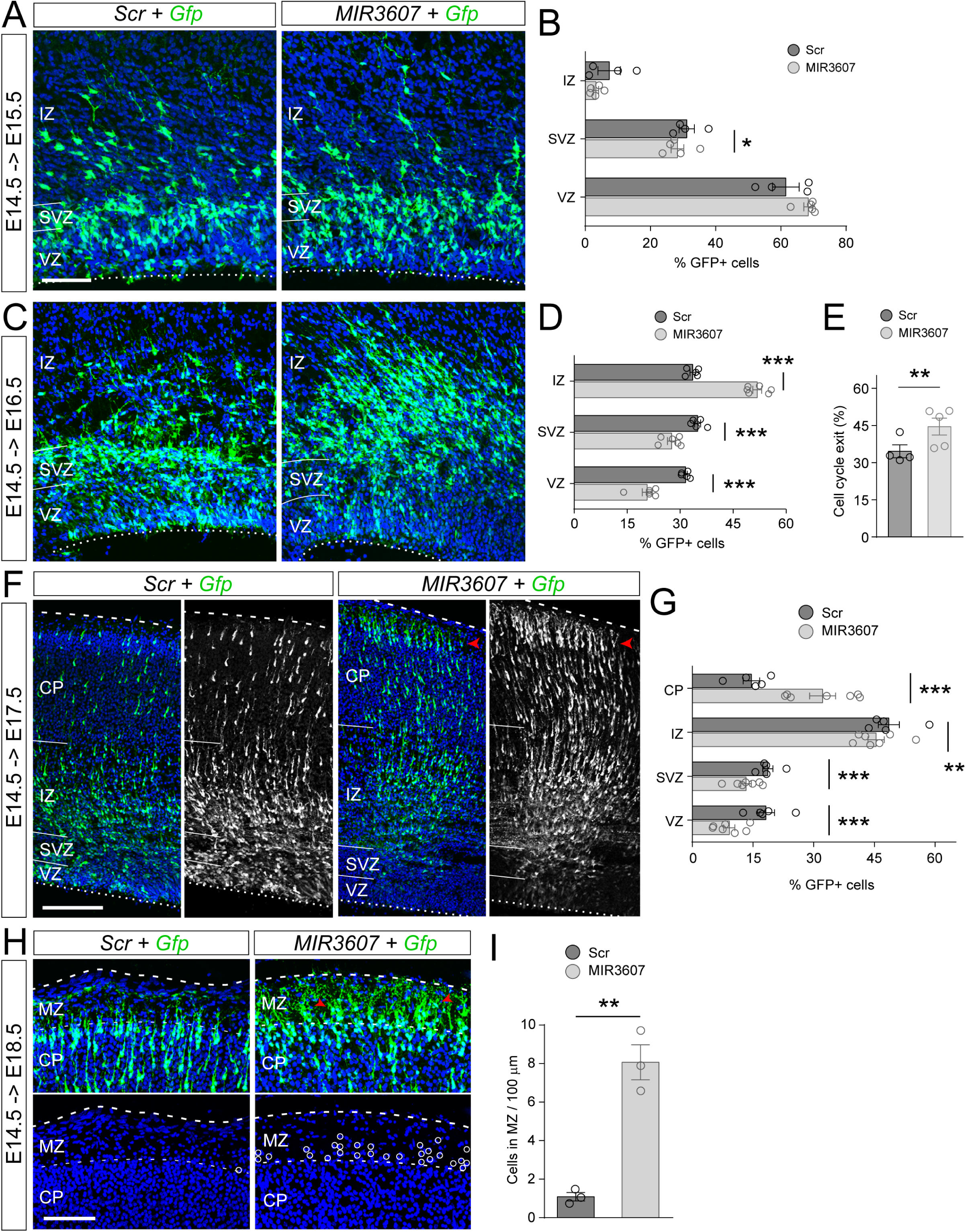
Defects in neurogenesis and neuron migration upon expression of *MIR3607* in mouse neocortex. (**A-D**) Sections through the parietal cortex of E15.5 (A) and E16.5 (C) mouse embryos electroporated at E14.5 with *Gfp* plus *Scrambled-* (*Scr*) or *MIR3607*-encoding plasmids, and quantifications of laminar distribution of GFP+ cells (B and D, respectively). Dotted lines indicate apical surface of the embryonic cortex. (**E**) Quantification of cell cycle exit rate at E16.5 for cells electroporated at E14.5 and receiving a single pulse of BrdU at E15.5. (**F,G**) Sections through the parietal cortex of E17.5 mouse embryos electroporated at E14.5 with the indicated plasmids (F), and quantifications of laminar distribution of GFP+ cells (G). Dotted lines indicate apical surface of the cortex, dashed lines indicate basal surface. (**H,I**) Detailed images of the marginal zone (MZ) and cortical plate (CP) of E18.5 mouse embryos electroporated at E14.5 with the indicated plasmids, showing high density of ectopic GFP+ cells within MZ in *MIR3607* embryos (H, open circles), and their quantification (I). Circles in plots indicate values for individual embryos. Histograms indicate mean ± SEM; *n* = 3-7 embryos per group; *X^2^* test (B,D,E,G), t-test (I); \**p*<0.05, \*\**p*<0.01, \*\*\**p*<0.001. Scale bars: 100µm (A,C,H), 200µm (F).

Next, we reasoned that if *MIR3607* expression increased the ratio of neurogenic over self-renewing divisions, too many neurons would be generated too early. This would in turn result in an increase of radially-migrating neurons in IZ at subsequent stages, as we observed at E16.5 (Fig. 2D), and also the subsequent arrival of these neurons to the CP ahead of control-electroporated neurons. In agreement with this notion, E17.5 embryos expressing *MIR3607* since E14.5 showed lower proportions of GFP+ cells in VZ, SVZ and IZ, and a two-fold increase of GFP+ cells in CP (Fig. 2F,G). In fact, in contrast to cells in control embryos, *MIR3607* expressing cells already started to settle and accumulate at the top of CP (Fig. 2F), where neurons end radial migration and start differentiating. These results seemed to indicate that the primary effect of *MIR3607* expression was to promote neurogenesis, which secondarily translated into the premature radial migration and settling of neurons in the CP. However, analyses one day later (E18.5 embryos electroporated at E14.5) revealed that in *MIR3607* embryos many GFP+ cells actually migrated past their arrival zone at the CP-MZ border, invading ectopically the MZ (Fig. 2H,I), which indicated a defective termination of radial migration.

Defects in neuron migration upon expression of *MIR3607* were confirmed at P5, after cortical lamination is complete (Fig. 3A). GFP+ neurons in miR-electroporated embryos were found in the CP (prospective layer 2/3 at this stage) and expressed the layer 2/3 marker Cux1, exactly like in control embryos, consistent with them maintaining a normal laminar fate (Fig. 3A-A’’). However, GFP+ neurons of *MIR3607* embryos occupied the top of CP, whereas in control-electroporated embryos they mostly occupied central positions within the CP (Fig. 3B,C). Considering that expression of *MIR3607* induced premature neurogenesis, according to the inside-out gradient of cortex development this should have resulted in neurons accumulating in deep positions within the CP, not superficial. However, our result was consistent with the overmigration defect observed previously at E18.5. In addition to defects in the CP, we also observed that many *MIR3607* neurons accumulated in the cortical white matter, forming subcortical ectopias (Fig. 3A). Marker analysis showed that the vast majority of these ectopic neurons expressed Cux1 (Fig. 3D), demonstrating retention of their normal fate for layer 2/3. Together, these observations confirmed that *MIR3607* expression altered radial neuron migration in the developing mouse cerebral cortex.

**Figure 3.**
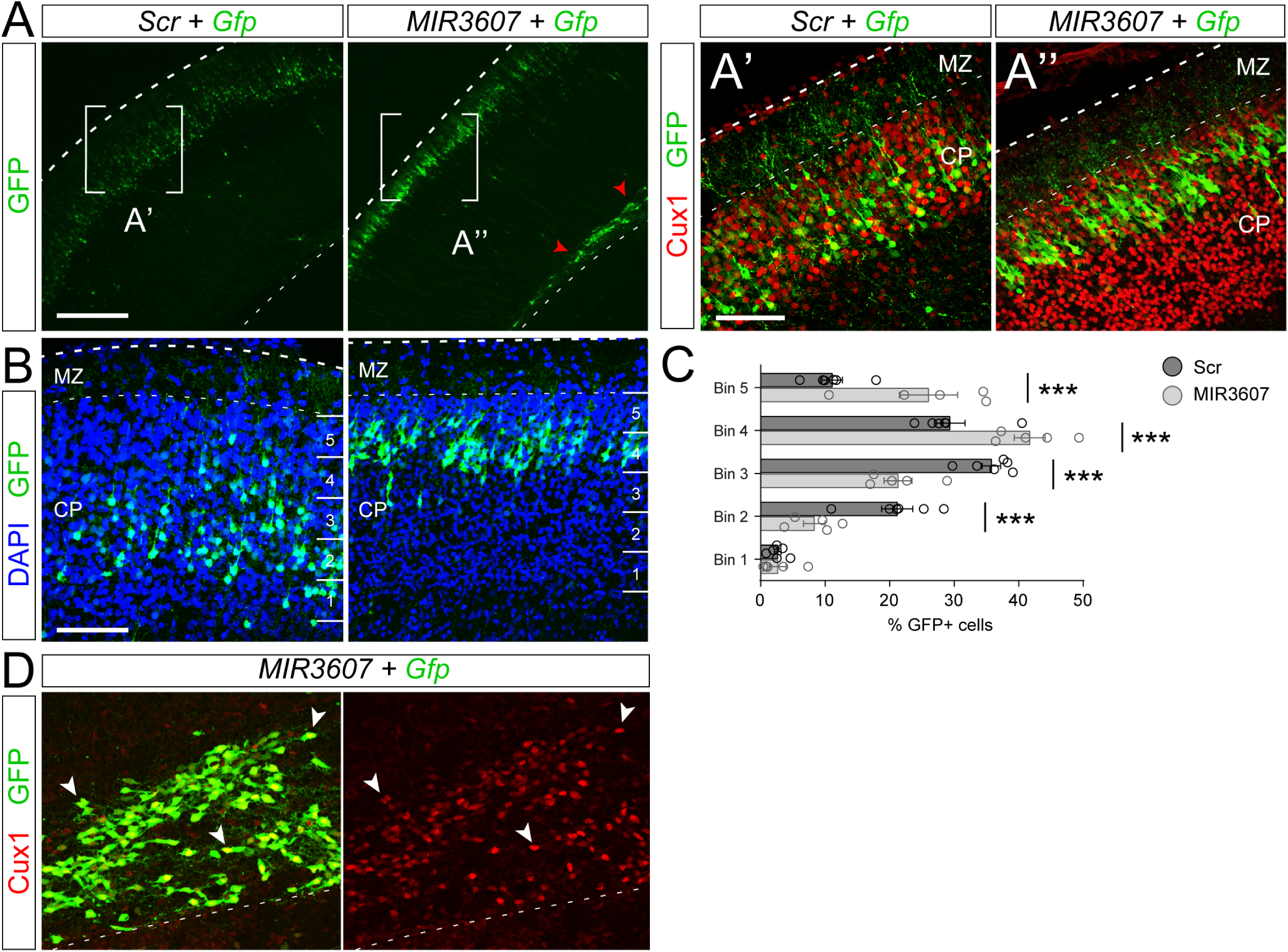
*MIR3607* expression causes long-term defects in neuron position. (**A-D**) Sections through the parietal cortex of P5 mouse pups electroporated at E14.5 with *Gfp* plus *Scrambled-* (*Scr*) or *MIR3607*-encoding plasmids and stained as indicated, and quantification of the binned distribution of GFP+ cells within the CP (B, C). (A’-A’’) are magnifications of the regions boxed in (A). (D) is a magnification of the white matter. All GFP+ cells in CP (normotopic) and white matter (ectopic) were positive for the upper layer marker Cux1 (red). Expression of *MIR3607* caused the overmigration and persistent presence of cortical neurons at the top of CP, and simultaneously the accumulation of ectopic neurons in the white matter (arrowheads in A and D), all of which maintained a correct upper layer fate. Circles in plots indicate values for individual embryos. Histogram indicates mean ± SEM; *n* = 5-6 embryos per group; *X^2^* test; \*\*\**p*<0.001. Scale bars: 300µm (A), 100µm (A’,A’’,B,D).

Given the effects of *MIR3607* on neurogenesis and neuron migration, we next enquired whether axonal growth was also affected. We electroporated E14.5 embryos to target progenitor cells producing layer 2/3 neurons, and then analyzed their growing axons across the Corpus Callosum (CC) at P5. Both in control and *in MIR3607* expressing embryos, GFP+ axons crossed the telencephalic midline at the level of the CC, and extended along the white matter of the contralateral hemisphere (**Supp Fig. S2A**). However, the density of GFP+ callosal axons extending along the white matter was lower in *MIR3607* embryos than in controls. Importantly, the deficit in callosal axons increased as these approached the midline, while after midline crossing it remained largely unchanged (**Supp Fig. S2B,C**). In the contralateral cortex, we further observed the invasion of axons from the white matter toward the CP in control embryos, which was virtually absent in *MIR3607* embryos (**Supp Fig. S2D**).

Taken together, our results demonstrated that expression of *MIR3607* affects development of the cerebral cortex at multiple levels. It induces premature neurogenesis, alters radial migration of neurons leading to defects in lamination and formation of white matter ectopias, and it impairs growth of cortical callosal axons, both in their initial navigation toward the midline and in their subsequent invasion of the contralateral grey matter.

### *MIR3607* drives amplification of Pax6+ progenitor cells

In the above analyses we found that expression of *MIR3607* at E14.5 in mouse cortical progenitor cells increased cell cycle exit and premature neurogenesis by E16.5, with a significant loss of cells in VZ and SVZ, and gain of IZ cells. However, one day earlier (E15.5) we found more cells in VZ and fewer in IZ and SVZ, suggesting that the immediate early effect might be the opposite: reduced neurogenesis and increased aRGC amplification. In agreement with this notion, BrdU labeling analyses revealed a 36% increase in BrdU-incorporating cycling progenitor cells in E15.5 embryos expressing *MIR3607* compared to control littermates (Fig. 4A,B), and a 65% reduction in cell cycle exit between E14.5 and E15.5 (Fig. 4B). This confirmed that the earliest changes immediately upon *MIR3607* expression are reduced neurogenesis and dramatic amplification of progenitor cells.

**Figure 4.**
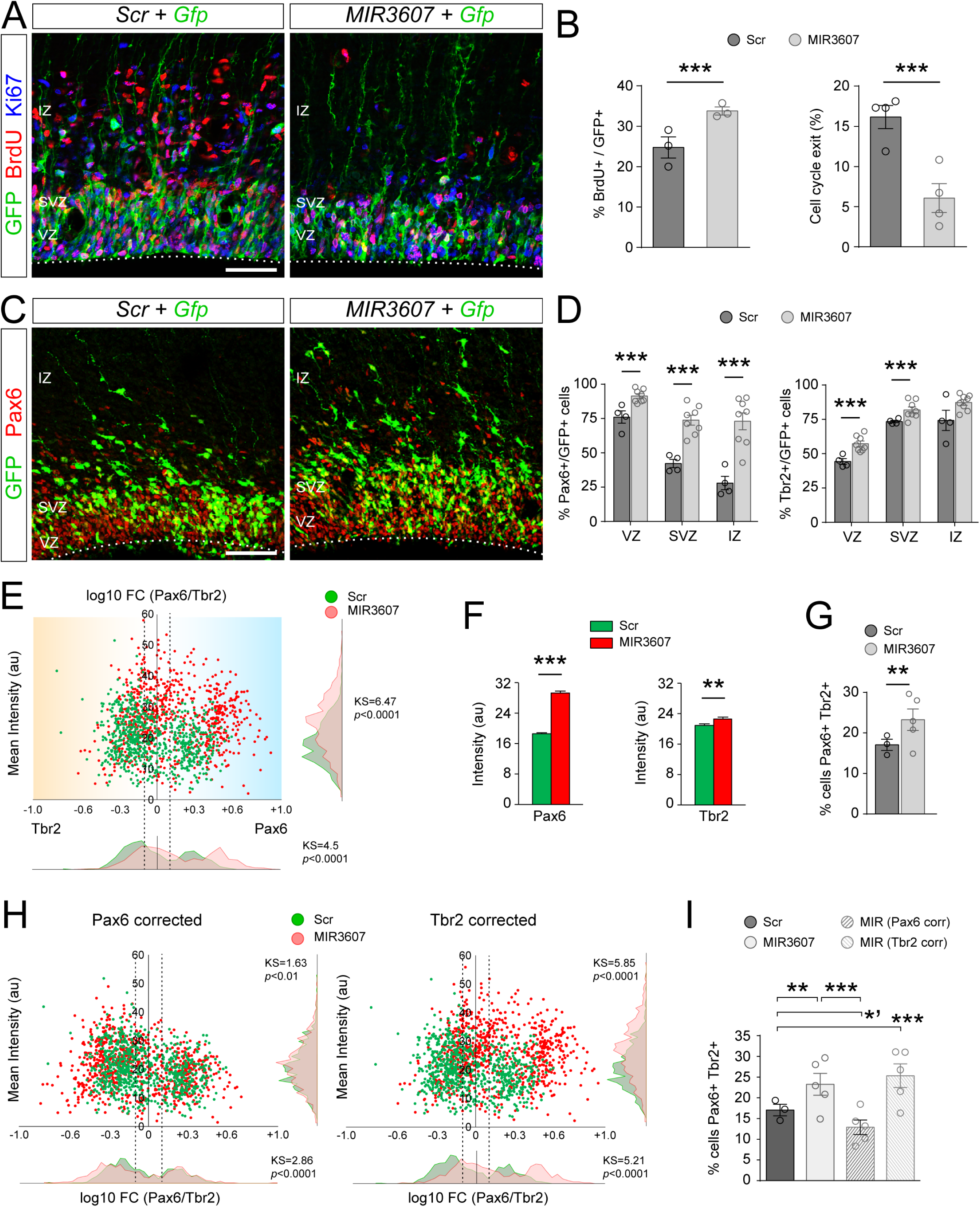
Amplification of Pax6+ progenitor cells immediately upon *MIR3607* expression. (**A,B**) Sections through the parietal cortex of E15.5 mouse embryos electroporated at E14.5 with *Gfp* plus *Scr*- or *MIR3607*-encoding plasmids stained as indicated, and quantifications of BrdU labelling index and cell cycle exit of GFP+ cells. (**C,D**) Distribution and abundance of Pax6+ and Tbr2+ cells in the parietal cortex of embryos electroporated as in (A). GFP+ cells in *MIR3607*-expressing embryos had greater BrdU intake, lower cell-cycle exit, and displayed a dramatic increase in frequency of Pax6 expression. (**E**) Scatter plots of ratio Pax6/Tbr2 expression level (log10 Fold Change) over their mean intensity (arbitrary units) in individual GFP+ cells as in (C), and frequency distribution plots for each of the two parameters, from *Scr-* and *miR*-expressing embryos. Dashed vertical lines delimit Pax6/Tbr2 co-expression (−0.1<log10FC<+0.1). *N* = 738 cells, 3 embryos for *Scr*; 790 cells, 5 embryos for *MIR3607*. (**F,G**) Average expression intensity of Pax6 and Tbr2 among individual cells (F), and proportion of cells co-expressing both factors (G), as defined in (E). (**H**) Scatter plots and frequency distribution plots as in (E), but where Pax6 or Tbr2 values of each *MIR3607* cell were corrected to the average *Scr* value as in (F). (**I**) Proportion of cells co-expressing Pax6 and Tbr2, as defined in (E), for the indicated conditions. The increased Pax6/Tbr2 co-expression in *MIR3607* cells was rescued *in silico* with Pax6 correction, and hence it was due to increased Pax6 intensity. Open circles in plots indicate values for individual embryos. Histograms indicate mean ± SEM; *n* = 3-8 embryos per group. *X^2^* test (B,D,G,I), t-test (F), Kolmogorov-Smirnoff test (E,H); *’*p*<0.02, \*\**p*<0.01, \*\*\**p*<0.001. Scale bars: 100µm.

The vast majority of mouse cortical progenitor cells in VZ are aRGCs, which characteristically express the transcription factor Pax6, whereas most progenitors in SVZ are IPCs, recognized by expression of the transcription factor Tbr2 (Englund et al., 2005; Gotz et al., 1998). To determine if the amplification of progenitor cells immediately upon *MIR3607* expression regarded aRGCs or IPCs, we next analyzed the expression of Pax6 and Tbr2 in E15.5 embryos. In control embryos, a majority (76%) of GFP+ cells in VZ were positive for Pax6, as expected (Fig. 4C,D). Pax6 was also expressed by 42% of SVZ and 28% of IZ cells. Tbr2 was detected mostly in SVZ and IZ cells (74%), and to a lesser extent in VZ (44%; Fig. 4D). In embryos expressing *MIR3607*, the proportion of Pax6+ cells increased in all three layers, most dramatically in SVZ and IZ (1.7- and 2.6-fold, respectively; Fig. 4D). Tbr2+ cell abundance also increased in VZ and SVZ upon expression of *MIR3607*, but much less than Pax6+ cells (Fig. 4D). While increased Pax6+ cells in VZ was consistent with greater abundance of aRGCs, it was puzzling to find such high frequency in SVZ and IZ, because these layers are normally populated by IPCs and newborn neurons, negative for Pax6 (Arai et al., 2011; Englund et al., 2005). In fact, the high frequencies of Pax6+ and Tbr2+ cells in *MIR3607* embryos were only compatible with a significant increase in the co-expression of these two proteins, which is one of the defining features of OSVZ progenitors in ferret and primates (Betizeau et al., 2013; Reillo & Borrell, 2012; Reillo et al., 2017). Analyses of staining intensity in individual cells demonstrated an average 60% increase in Pax6 levels upon *MIR3607* expression, with only an 8% increase in Tbr2 levels (Fig. 4E,F). This selective change resulted in a significant increase in the abundance of cells co-expressing Pax6 and Tbr2 (Fig. 4E,G), largely attributable to basal cells in SVZ (**Supp Fig. S3**). *In silico* analyses revealed that the observed increase in Pax6 levels, but not Tbr2, was sufficient to explain the differences between control and *MIR3607* expressing embryos in Pax6-Tbr2 co-expression (Fig. 4H,I).

### *MIR3607* activates signaling pathways driving progenitor proliferation

To elucidate the mechanism of action of *MIR3607* leading to the changes in cortical progenitor cells reported above, we next investigated the impact of *MIR3607* expression at the transcriptomic level. We electroporated *in utero* the cerebral cortex of E14.5 embryos with plasmids encoding *MIR3607* or *Scrambled* plus GFP, and 24h later RNA-sequencing (RNA-seq) was performed on FACS-purified GFP positive cells (Fig. 5A). We identified 173 genes differentially expressed (DEGs) in cortical progenitors upon *MIR3607* expression (FDR<0.01; Fig. 5B, **Supp Table S1**). A majority of DEGs were downregulated (58%), as expected from the action of a miRNA (Fig. 5B). Of the 76 genes in the mouse genome computationally predicted to be direct targets of *MIR3607*, 63 were expressed in our samples, and 9 of them were DEGs (Fig. 5C). The great majority of those DEGs were downregulated (8 out of 9), again as predicted from the action of a miRNA: Dnm3, Opcml, Pde4d, Tmem169, Cnr1, Bsn, Apc and Rnf38 (Fig. 5C). Several functional enrichment analyses were performed to capture biological information on DEGs. Gene Ontology (GO) analysis highlighted the Wnt signaling pathway and axon development as having the highest enrichment (Fig. 5D). Similarly, functional grouping of gene networks highlighted Wnt signaling pathway, neuroblast proliferation, regulation of neural precursor cell proliferation and L1CAM interactions (Fig. 5E). L1-CAM interactions are relevant for axon development, so these results were consistent with the observed deficient growth of callosal axons in P5 mice expressing *MIR3607* (**Supp Fig. S2**). Functional annotation clustering analysis highlighted again, as top ranked, a cluster topped by the terms Wnt signaling pathway, lateral plasma membrane and signaling pathways regulating pluripotency of stem cells (Fig. 5F). Altogether, these analyses revealed a prominent role of *MIR3607* in the regulation of Wnt/β-catenin signalling pathway, proliferative activity, axon development and lateral plasma membrane (Fig. 5D-F; **Supp Table S1**). All these biological functions closely matched the developmental processes that we found altered in our above phenotypic analyses of the developing cortex upon expression of *MIR3607* at E14.5 (Figs. 2-4). Indeed, *MIR3607* expression in cortical progenitor cells led to decreased expression of Adenomatous Polyposis Coli (APC), a key negative modulator of the canonical Wnt/β-catenin signalling pathway, and concomitantly to increased levels of Fgfr3, Fzd8, Ctnn1a and Ctnn1b transcription (Fig. 5B,C,E-H). Consistent with the GO analysis, gene set enrichment analyses (GSEA) (Subramanian et al., 2005) further confirmed a strong modulation by *MIR3607* of genes regulating Wnt/β-catenin signalling, the apical junctional complex and cell division in cortical progenitors (Fig. 5I-K, **Supp Fig. S5, Supp Table S1**). In summary, our transcriptomic analyses revealed that *MIR3607* expression causes dramatic changes in the expression levels of genes participating in biological functions and signaling pathways that are key for the amplification and delamination of cortical progenitor cells.

**Figure 5.**
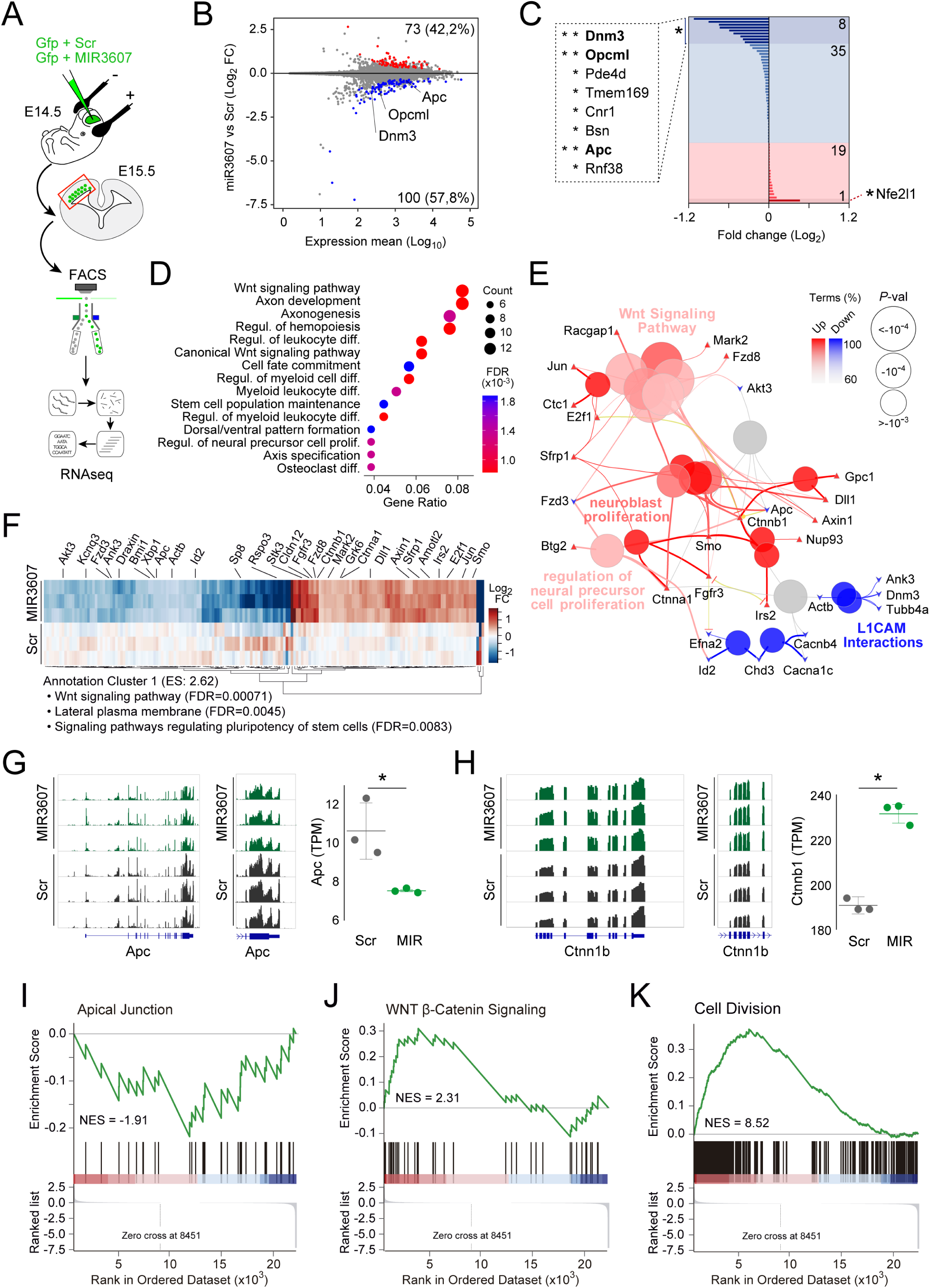
*MIR3607* promotes cell proliferation and activates Wnt signaling. (**A**) Schema of experimental design: E14.5 mouse embryos were electroporated with *Gfp* plus *Scr*- or *MIR3607*-encoding plasmids, at E15.5 their brains were microdissected and dissociated, cells expressing high levels of GFP were purified by FACS sorting, and their pooled RNA expression profiles were analyzed by RNAseq. (**B**) Plot of Fold Change in gene expression (MAP estimate) in *MIR3607-* versus *Scr*-electroporated cells, over the gene’s average expression level. Differentially Expressed Genes (DEGs; Adj. P < 0.01) are in red (upregulated, 73 genes) and blue (downregulated, 100 genes). Top three DEGs among *MIR3607*-predicted targets are named. (**C**) Fold change of 63 predicted *MIR3607* targets detected. DEGs are named (*Adj. *p*<0.1; **Adj. *p*<0.01); 8 genes were downregulated, and only one upregulated. (**D**) Functional enrichment analysis on DEGs, showing most significant enriched GO terms. Ontology: Biological Process. (**E**) Functionally grouped network based on functional enrichment analysis of DEGs. Size of nodes indicates statistical significance of the terms (Bonferrroni step down corrected P-values). Percentage of up- and down-regulated DEGs for each term is indicated by the color of the nodes on the network. (**F**) Heatmap of DEGs highlighting genes associated to the top ranked Annotation Cluster from Functional Annotation Clustering analysis (DAVID 6.8). The top three terms within this cluster and their statistical significance (FDR) are indicated. (**G,H**) Visualization of normalized coverage tracks from RNAseq data for whole transcript (left, middle), and expression levels (right) for *Apc* (G) and *Ctnn1b* (H) in *Scr*- and *MIR3607*-expressing cells. TPM, transcripts per million; *n* = 3 replicates per condition; Student two sample t-test; \**p*<0.05. (**I-K**) Enrichment plots from GSEA for MSigDB Hallmark Apical Junction (NES = −1.91; *p*=0.01; Adj. *p*=0.021), WNT β-catenin Signaling (NES = 2.31; *p*=0.002; Adj. *p*=0.006) and GO term Cell Division GO:0051301 (NES = 8.52; *p*=0.002; Adj. *p*=0.008).

### Massive delamination and amplification of aRGCs in the early cortex by *MIR3607*

Progenitor cell amplification and self-renewal are singularly important at early stages of cortical development, prior to and at the onset of neurogenesis, so we next investigated the effects of expressing *MIR3607* in the early embryonic mouse cerebral cortex. To this end, we electroporated *MIR3607*-encoding plasmids into the cortical primordium of mouse embryos at E12.5. To investigate the immediate early effects we analyzed the cortex of electroporated embryos at E13.5, which revealed severe alterations in the organization of progenitor cells (Fig. 6). BrdU labeling experiments revealed that cells in S-phase were perfectly aligned in the basal aspect of the VZ in control-electroporated and non-electroporated cortices (Fig. 6A,B). In contrast, expression of *MIR3607* caused the spreading of BrdU-incorporating cells over the entire thickness of the VZ (Fig. 6B). Remarkably, BrdU+ cells were frequently arranged in circles, resembling rosettes (Fig. 6A), which suggested increased rate of cell proliferation and amplification. This disorganization was paralleled by a severe disruption of the apical adherens junction belt, as identified by Par3. In the area of cortex expressing *MIR3607*, Par3 was completely absent from the apical surface, instead forming small closed domains within the cortical parenchyma, in the lumen of rosettes (Fig. 6C). Accordingly, the typical band of PH3+ mitotic cells in the apical side of the normal VZ was not recognizable in *MIR3607* embryos, which instead displayed an excess of abventricular mitoses across the VZ thickness (Fig. 6D). The distribution of Pax6+ and Tbr2+ cells was similarly disrupted in agreement with a highly disorganized VZ (Fig. 6E,F). Importantly, the disorganization of Pax6+ and Tbr2 cells was consistent with these being constituent parts of rosettes: an inner mass of Pax6+ cells adjacent to Par3 forming the core of rosettes, and this being surrounded by Tbr2+ cells (Fig. 6C,E,F).

**Figure 6.**
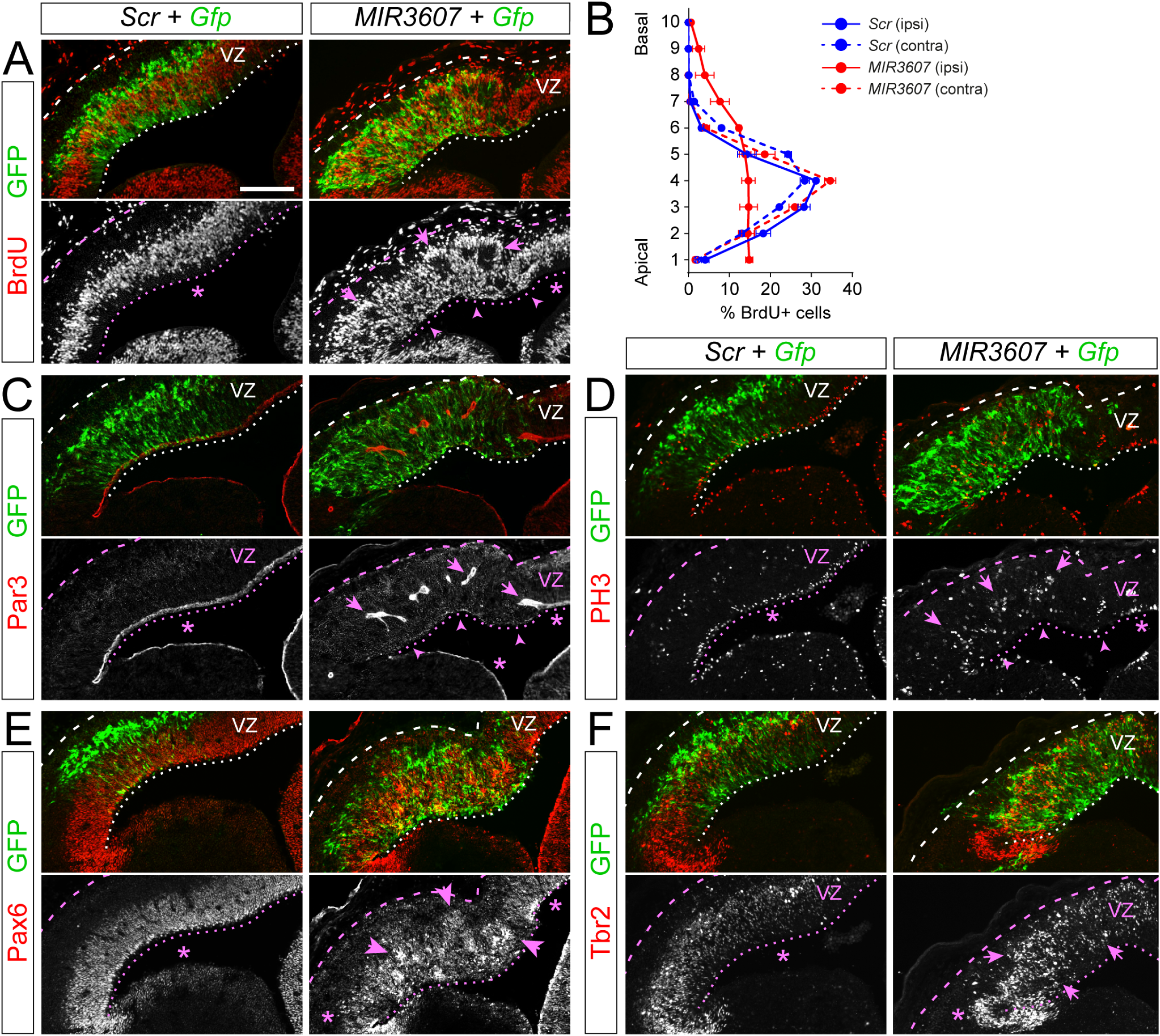
*MIR3607* drives massive delamination of apical progenitors and formation of rosettes. (**A,C-F**) Sections through the parietal cortex of mouse embryos electroporated at E12.5 with the indicated plasmids, analysed at E13.5 and stained as indicated. Expression of *MIR3607* caused a severe disruption of the laminar organization of cycling progenitors (A), the Par3+ apical adherens junction belt (C) and the distribution of PH3 mitoses (D), Pax6+ RGCs (E) and Tbr2+ IPCs (F) (arrows). Arrowheads indicate areas with disrupted apical side of VZ. Arrows indicate rosettes of progenitor cells delaminated into the cortical parenchyma. Asterisks indicate areas with normal organization. (**B**) Quantification of binned distribution of BrdU+ cells across the cortical thickness. Data is from electroporated hemispheres (solid lines) and from non-electroporated, contralateral hemispheres (dashed lines). The typical accumulation of BrdU-incorporating cells in the basal side of the VZ (bins 3-5) was observed in *Scr*-electroporated (solid blue line) and contralateral hemispheres (dashed lines), but severely disturbed in *MIR3607* electroporated cortices (solid red line). Plots show mean ± SEM; *n* = 2,673 cells ipsi, 3,583 cells contra, 2 embryos, *Scr*; 5,150 cells ipsi, 5,485 cells contra, 4 embryos, *MIR3607*. Scale bar: 100µm (all panels).

The general disorganization of germinal layers in *MIR3607*-electroporated embryos persisted at later stages (E15.5), becoming even more dramatic. Cycling BrdU+ progenitor cells failed to remain within the basal side of VZ, typical of control embryos, but were wide spread through the SVZ and IZ (Fig. 7A). Stains against Par3 demonstrated the persistent absence of the apical adherens junction belt in VZ of *MIR3607* expressing embryos, and their continued presence as small circular structures at basal positions within the cortical parenchyma (Fig. 7B). Pax6 stains showed that in *MIR3607* expressing embryos aRGCs no longer formed a compact VZ as in controls, but spread from the apical VZ surface to the IZ (Fig. 7C). These stains also revealed clusters of Pax6+ cells located basally, coincident with the location of rosettes. This massive disorganization of the VZ with delamination of Pax6+ aRGCs also affected IPCs, identified with Tbr2 stains. Tbr2+ cells were also distributed ectopically in miR-expressing embryos, where they extended apically through the thickness of VZ to its apical surface, and basally through SVZ and IZ, where they formed distinct circular clusters (Fig. 7D). At the level of cell divisions, expression of *MIR3607* led to a 65% loss of PH3+ apical mitoses alongside a 3-fold increase in basal mitoses. These no longer aligned in a distinct SVZ, but spread basally through the IZ, occasionally forming small clusters (Fig. 7E,F). Importantly, the combined abundance of apical and basal mitoses was 48% greater in *MIR3607*-expressing cortices than in controls (Fig. 7G), demonstrating increased progenitor cell proliferation. Overall, rosettes in the cortex of E15.5 *MIR3607* embryos displayed the same features as observed in early embryos: a Par3+ lumen, an apical layer of Pax6+ cells, and a surrounding basal layer of Tbr2+ cells. Detailed examination of GFP+ cells forming the core of rosettes confirmed their aRGC morphology, with distinct apical and basal processes radially aligned, and mitosis at the apical surface (Fig. 7H).

**Figure 7.**
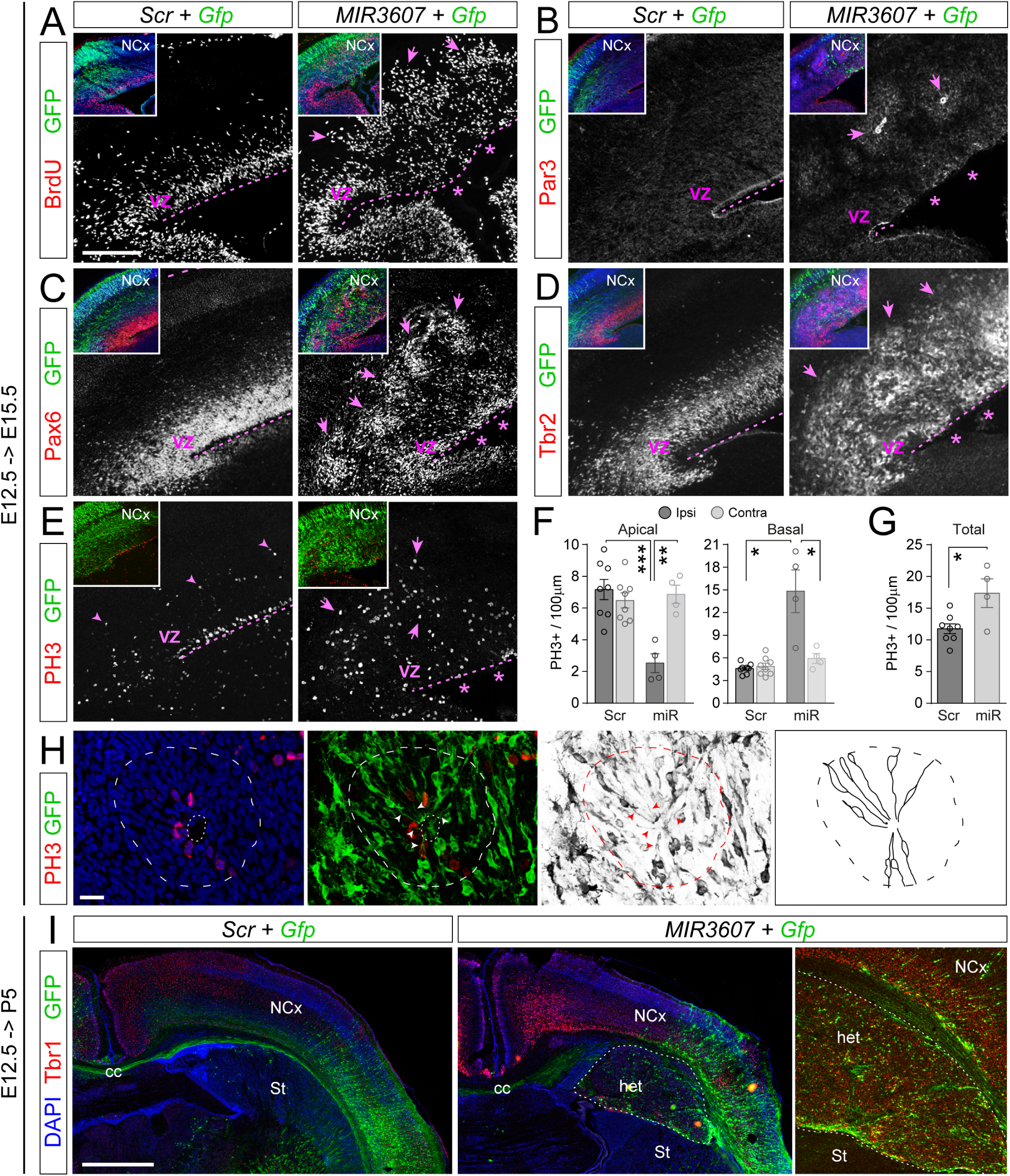
Severe disruption of apical junctions and formation of rosettes by *MIR3607* lead to subcortical heterotopia. (**A-D**) Sections through the parietal cortex of mouse embryos electroporated at E12.5 with the indicated plasmids, analysed at E15.5 and stained with the indicated markers. Expression of *MIR3607* disrupted severely the laminar organization of VZ and the Par3+ apical adherens junction belt (asterisks). Arrows indicate rosettes of cycling (BrdU+), Pax6+ and Tbr2+ progenitor cells, containing a Par3+ lumen, delaminated into the cortical parenchyma. Insets in each image are low magnifications showing DAPI (blue) and GFP expression in the same area. (**E-G**) Distribution and quantification of mitotic cells in E15.5 embryos electroporated at E12.5 with the indicated plasmids. Histograms indicate mean ± SEM for the electroporated hemisphere (Ipsi) and the non-electroporated, contralateral hemisphere (Contra). Circles in plots indicate values for individual embryos. Expression of *MIR3607* severely decreased apical mitoses (asterisks) and increased basal mitoses (arrows), with an overall increase in mitotic density (G). *N* = 4-8 sections from 2-4 embryos per group; t-test; \**p*<0.05, \*\**p*<0.01, \*\*\**p*<0.001. (H) Detail of a rosette (dashed line) in the cerebral cortex of an E15.5 embryo electroporated with *MIR3607* at E12.5, showing apical PH3 mitoses in the inner lumen (dotted line) surrounded by the apical processes of GFP+ aRGCs (arrowheads). Right panel shows line reconstructions of GFP+ cells within the rosette, demonstrating typical aRGC morphology with apical and basal processes radiating from the inner lumen. (I) Sections through the parietal cortex of mouse embryos electroporated at E12.5 with the indicated plasmids, analysed at postnatal day (P) 5, and stained for GFP and the deep-layer marker Tbr1. Note the mass of Tbr1+ neurons in the heterotopia (het) between the normal neocortex (NCx) and striatum (St) of the brain expressing *MIR3607*. cc, corpus callosum. Scale bars: 200µm (A-E), 10µm (H), 1 mm (I).

We finally investigated the long-term consequences of *MIR3607* expression and this highly disorganized cortical development. Examination of P5 mouse pups electroporated at E12.5 revealed the formation of subcortical heterotopia underneath the electroporation site (Fig. 7I). The size of this heterotopia varied between animals, but was never observed in *Scramble*-electroporated control mice. Heterotopias were largely composed by cells positive for Tbr1, a marker of deep layer neurons, whereas a majority of cells were negative for GFP, indicating that the mechanism of emergence of this phenotype had a significant non-cell autonomous component (Fig. 7I).

In summary, the formation of rosettes seemed to involve the massive delamination of aRGCs followed by IPCs, with the corresponding loss of a distinctive VZ. This demonstrated that the effects of *MIR3607* expression in the early developing cortex included disruption of the neuroepithelial organization of germinal layers, and a very substantial increase in progenitor cell proliferation. These effects were fully consistent with our previous functional analyses of transcriptomic changes in these embryos, overall highlighting effects on neural progenitor cell proliferation, cell division, lateral plasma membrane and apical junction (Fig. 5), altogether leading to the formation of proliferative rosettes.

### *MIR3607* promotes early cortical progenitor amplification and formation of rosettes by de-repression of βCatenin signaling

The effects of expressing *MIR3607* in the embryonic mouse cerebral cortex, causing a massive amplification of cortical progenitors and disturbance of VZ integrity, resembled the effect produced by overactivation of the Wnt/βCatenin signaling pathway (Chenn & Walsh, 2002; Chenn & Walsh, 2003; Herrera et al., 2014; Poschl et al., 2013; Woodhead et al., 2006; Wrobel et al., 2007). Accordingly, our transcriptional profiling experiments revealed the strong and preferential activation of this pathway at the transcriptional level in the developing mouse cerebral cortex upon *MIR3607* expression (Fig. 5). We confirmed these transcriptomic results at the protein level, by immunostains against activated βCatenin, which revealed a two-fold increase in the VZ of *MIR3607* expressing mouse embryos 24hr after electroporation (Fig. 8A,B). This further supported that the effects of *MIR3607* on cortical progenitor amplification and rosette formation might be caused by the overactivation of βCatenin signaling. To investigate this possibility, we electroporated E12.5 embryos with a constitutively active form of βCatenin (Δ90-βCat). One day after electroporation (E13.5), the organization of the cerebral cortex and its VZ were severely disrupted, with multiple proliferative rosettes and massive disorganization of cycling progenitor cells (Fig. 8C,D), phenocopying the effects of *MIR3607* expression but at an even greater level of disruption. This further supported the notion that the alterations in cortical progenitor amplification and delamination caused by *MIR3607* were mediated by the overactivation of βCatenin signaling.

**Figure 8.**
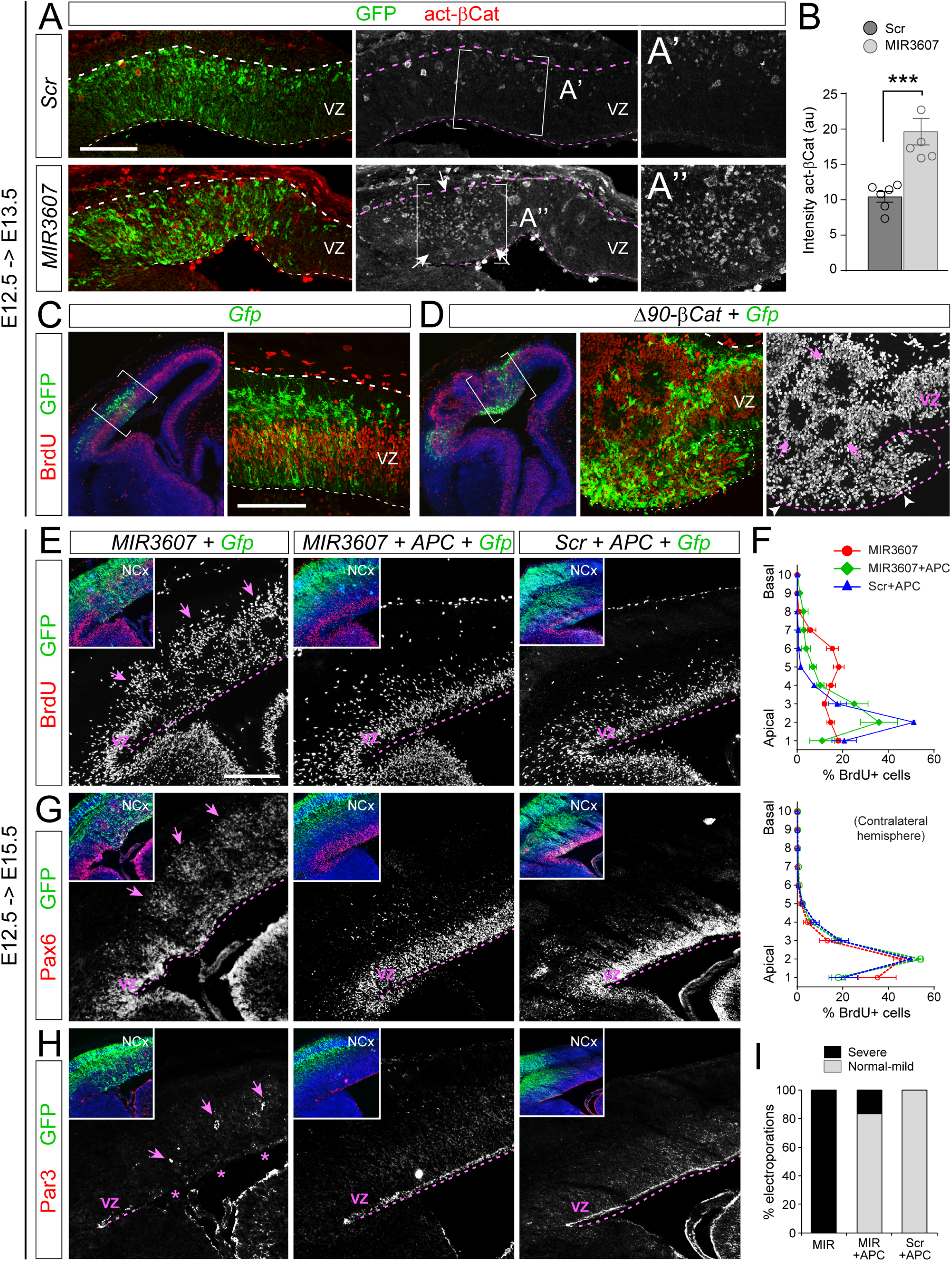
*MIR3607* promotes early progenitor amplification and formation of rosettes by de-repression of βCatenin signaling. (**A,B**) Sections through the parietal cortex of E13.5 mouse embryos electroporated at E12.5 with the indicated plasmids and stained against activated βCatenin, and quantification of signal intensity. (A’,A’’) are high magnifications of the corresponding areas boxed in (A). Histogram indicates mean ± SEM; circles indicate values for individual embryos; *n* = 5-6 embryos per group; t-test; \*\*\**p*<0.001. (**C,D**) Parietal cortex of E13.5 mouse embryos electroporated at E12.5 with the indicated plasmids. Expression of constitutively active βCatenin very severely disrupted the integrity of the VZ (white arrowheads), with massive amplification and delamination of cycling (BrdU+) progenitor cells and formation of rosettes (pink arrows). (**E-H**) Sections through the parietal cortex of E15.5 mouse embryos electroporated at E12.5 with the indicated plasmid combinations and stained as indicated (E,G,H), and binned distribution of BrdU+ cells across the cortical thickness (F). In (F), top graph is data from electroporated, ipsilateral hemispheres; bottom is data from non-electroporated, contralateral hemispheres. Asterisks indicate the absence of the Par3+ apical adherens junction belt; arrows indicate rosettes. The severe disruptions caused by *MIR3607* expression were rescued in embryos co-expressing *MIR3607* and APC, and absent in embryos expressing *Scr*+APC. (**I**) Quantification of embryos with germinal layer disturbance. Scale bars: 100µm (A-D), 200µm (E,G,H).

*MIR3607-5p* has been shown to target the 3’UTR of *APC*, a key repressor of βCatenin signaling, blocking its expression in a human lung cancer cell line (Lin et al., 2017). Moreover, our transcriptomic analyses showed a significant decrease of *APC* mRNA levels in mouse cortical progenitor cells expressing *MIR3607* (Fig. 5F,G). This suggested that *MIR3607* may activate βCatenin signaling indirectly, via repressing APC expression. If this was the case, the deleterious effect of *MIR3607* expression on the embryonic cerebral cortex should be rescued by additionally expressing APC. We tested this possibility by expressing APC alongside *MIR3607* in the E12.5 mouse cortex, and analyzed the effects at E15.5. As shown previously, expression of *MIR3607* alone caused a very severe disorganization of germinal zones in the developing cerebral cortex. Cycling progenitor cells, identified by BrdU incorporation, were distributed through the thickness of the VZ and spread basally to the IZ, forming conspicuous rosettes (Fig. 8E,F). The formation of these rosettes involved the severe disruption of Par3+ adherens junctions and the delamination of Pax6+ cells from VZ to SVZ and IZ, where they formed proliferative rosettes (Fig. 8E-H). Remarkably, co-electroporation of *MIR3607* with APC completely rescued all these defects in a majority of embryos, whereas expression of APC with a control, scrambled miRNA sequence, had no effect on cortical progenitor cells nor on the normal organization of germinal layers (Fig. 8E-I). Together, these results demonstrated that expression of *MIR3607* in the embryonic mouse cerebral cortex reduces the levels of *APC* expression, which leads to an abnormal accumulation of activated βCatenin and the overactivation of the canonical Wnt pathway, causing the overproliferation of cortical progenitor cells and their massive delamination from the VZ forming proliferative rosettes.

## Discussion

### Gene regulation and secondary reduction of cortex size during evolution

The mammalian cerebral cortex expanded dramatically during evolution. This expansion was very prominent in humans and great apes, but it took place in all major clades, including carnivores, ungulates and cetaceans. However, according to evolutionary modeling, in some clades this process was reversed during subsequent evolution, thereby undergoing a secondary reduction of brain size (Kelava et al., 2013). This has been shown for species with a small and smooth cortex, like small world monkeys such as marmoset, and rodents like mice, where the Capibara remains an exception as the only extant rodent with a large and folded brain (Garcia-Moreno et al., 2012; Kelava et al., 2013; Kelava et al., 2012). This secondary reduction of brain size is the result of variations during their embryonic development, which in fact recapitulate differences in evolution between species. For example, in species with a large and folded cerebral cortex like human, macaque monkey, sheep or ferret, its embryonic development involves a large abundance of neural stem and progenitor cells, particularly bRGCs, which coalesce in thick and complex subventricular germinal zones (ISVZ and OSVZ) (Betizeau et al., 2013; Fietz et al., 2010; Hansen et al., 2010; Reillo & Borrell, 2012; Reillo et al., 2011; Smart et al., 2002). In contrast, in species with a smaller and smooth cortex, like mouse or marmoset monkey, embryonic cortical development involves much fewer of those progenitor cells, with bRGCs being a small proportion of cortical progenitors, and frequently a simpler subventricular zone (Englund et al., 2005; Garcia-Moreno et al., 2012; Kelava et al., 2012; Wang et al., 2011). This circumstance provides a unique opportunity to test experimentally whether developmental features specific to large-brained species are important for the expansion of their cerebral cortex, by introducing them into small-brained species like mouse or marmoset. This strategy has been followed previously to demonstrate the effect on cerebral cortex size of decreased developmental apoptosis (Depaepe et al., 2005; Haydar et al., 1999; Kingsbury et al., 2003), increased neural stem cell proliferation (Chenn & Walsh, 2002; Nonaka-Kinoshita et al., 2013) and increased basal progenitor abundance (Stahl et al., 2013).

At the genetic level, an increasing number of transcriptomic studies have identified protein-coding genes expressed in germinal layers of the developing cerebral cortex that are either specific to large-brained species, or that are expressed at different levels according to brain size or depending on progenitor cell type (Fiddes et al., 2018; Fietz et al., 2012; Florio et al., 2015; Florio et al., 2017; Florio et al., 2018; Suzuki et al., 2018). Most importantly, functional testing of some of these genes demonstrates their relevance in the emergence or acquisition of features typical of species with large brains. This is the case for genes expressed in the developing cortex of both mouse and human, but at different levels: the mouse cerebral cortex undergoes significant expansion and/or folding upon overexpression of *Pax6*, *Hopx*, or *Fgf2*, or downregulation of *Trnp1* or *Flrt1/3* (Del Toro et al., 2017; Rash et al., 2013; Stahl et al., 2013; Vaid et al., 2018; Wong et al., 2015). A similar case is for primate- or human-specific genes like *ARHGAP11B*, *TMEM14B* and *NOTCH2NL*, which drive expansion of the cortical progenitor cell pool when expressed in the developing mouse cortex (Fiddes et al., 2018; Florio et al., 2015; Florio et al., 2018; Liu et al., 2017; Suzuki et al., 2018).

The number of human-specific genes, newly emerged during recent evolution, is relatively very small (Florio et al., 2018), whereas the number of conserved genes that are expressed in the developing cortex at different levels across phylogeny is much larger (Fietz et al., 2012; Florio et al., 2017). Hence, the key question now emerging is how the expression levels of such highly conserved genes became regulated so differently during evolution. Candidate mechanisms contributing to modify gene expression levels across species include gain, loss or modification of regulatory elements (promoters, enhancers), differential expression of transcriptionally-relevant protein-coding genes (transcription factors) and of non-coding genes (microRNAs and other small RNAs). In spite of the well-demonstrated power of miRNAs to modulate the expression of protein-coding genes (at the level of transcription or translation), they have received surprisingly little attention in the context of brain evolution and expansion (Arcila et al., 2014; Nowakowski et al., 2018). Here we have investigated this largely unexplored possibility focusing on *MIR3607*, a very attractive candidate miRNA for three main reasons: first, it has a highly conserved sequence across mammals with very different brain sizes, including human, ferret and mouse. Second, it is computationally predicted to target a large set of genes functionally related to early neural development, including neurogenesis and axon formation. Third, it is expressed in the developing cerebral cortex of large-brained species like human and ferret, but not of the small-brained mouse. Our results in mouse demonstrate that expression of *MIR3607* in the embryonic cerebral cortex is in itself sufficient to strongly promote the proliferation of cortical progenitor cells and expansion of their pool, and to drive their delamination from the VZ to basal positions, while retaining Pax6 expression. These are features greatly enhanced in the developing cerebral cortex of human, macaque and ferret, linked to the abundant formation of bRGCs (Fietz et al., 2010; Hansen et al., 2010; Reillo & Borrell, 2012; Reillo et al., 2011; Smart et al., 2002), and contribute critically to the large size of these cortices (Fernandez et al., 2016; Nonaka-Kinoshita et al., 2013; Reillo et al., 2011). In agreement with this line of reasoning, when we overexpressed *MIR3607* in human cerebral organoids, the VZ was fragmented into an increased number of closed proliferative ventricles, resembling delaminated rosettes (**Supp Fig. S6A,B,E**). Moreover, the length of the apical surface of these electroporated ventricles, decorated with the adherens junction protein Par3, was much greater in *MIR3607*-electroporated organoids than in controls (**Supp Fig. S6C,D,F**), consistent with the VZ progenitor cell amplification that we found in mouse embryos. Hence, in cerebral human organoids expression of *MIR3607* also increased proliferation and delamination of VZ. In summary, our results suggested that expression of *MIR3607* in the developing cortex may have been secondarily lost during rodent evolution as a mechanism to contribute to reduce cerebral cortex size. Testing this notion will require further investigations on the expression of mature miRNAs in the developing cerebral cortex, covering additional clades and brain phenotypic diversity across mammalian phylogeny.

### Post-transcriptional influence of *MIR3607*

An important concept strengthened by our current study is the relevance of post-transcriptional regulatory mechanisms on the evolution of brain development. MiRNAs are major components of this level of gene expression regulation, mostly controlling mRNA abundance (driving its degradation) or its translation into protein (blocking it) (Gebert & MacRae, 2019). The first mechanism has been extensively investigated in the Notch signaling pathway, where *miR-9* drives *Hes1* mRNA for degradation. The expression of *miR-9* and its effect on *Hes1* levels is key to modulate the oscillatory versus sustained levels of this Notch effector protein (Bonev et al., 2012), which in turn defines the fate of neural progenitor cells as self-amplifying or differentiating into neurons or glia (Goodfellow et al., 2014; Imayoshi et al., 2013; Shimojo et al., 2008). Tight regulation of translation is also common in the developing brain, where retention at high abundance of untranslated mRNA species is used as an efficient switch mechanism in protein activity. The transcription factors Pax6 and Tbr2 play key roles in cortical development by defining the identity of aRGCs and IPCs, respectively (Sessa et al., 2008). IPCs are born from Pax6+ aRGCs, but co-expression of Pax6 and Tbr2 at the protein level are near mutually-exclusive in mouse (Arai et al., 2011), which strongly suggests the involvement of both a very rapid degradation of Pax6 and translation of Tbr2. This is supported by the abundant presence of aRGCs within VZ that express high levels of Tbr2 mRNA, but not protein, in mouse as well as in ferret (de Juan Romero et al., 2015; Sessa et al., 2008). Because of the abundant accumulation of Tbr2 mRNA, this can be quickly and massively translated in IPCs immediately following their birth from aRGCs. This is in turn essential for the rapid reduction of Pax6 protein levels and consequent blockade of the aRGC fate (Sansom et al., 2009; Sessa et al., 2008). Interestingly, *miR-7* has been shown to inhibit Pax6 protein expression without altering its mRNA levels (Needhamsen et al., 2014; Zhang et al., 2018). Moreover, the 3’UTR of Pax6 bears a functional *miR-7* target site only in primates, suggesting that this interaction emerged during evolution linked to brain expansion (Needhamsen et al., 2014). Indeed, knocking down *miR-7* reduces the transition from aRGCs to IPs, and results in microcephaly-like brain defects (Pollock et al., 2014). Importantly, Pax6 and Tbr2 protein are very frequently co-expressed within basal progenitors of the developing cortex of macaque and ferret, essentially bRGCs (Betizeau et al., 2013; Reillo & Borrell, 2012), suggesting the absence, or override, of post-transcriptional regulation in these two particular genes. This in turn suggests the potential relevance of this absence in the acquisition of developmental features important for cortical expansion, including aRGC delamination and acquisition of bRGC fates.

We have shown that expression of *MIR3607* in the embryonic mouse cortex increases the frequency of cells co-expressing Tbr2 and Pax6 protein in VZ and, especially, in SVZ. This is particularly dramatic for Pax6, which we also found is expressed at much higher levels relative to Tbr2 in *MIR3607* expressing embryos. Intriguingly, neither *Pax6* nor *Tbr2* are computationally predicted targets of *MIR3607*, which is in agreement with their mRNA not being differentially expressed in *MIR3607* embryos, as found in our transcriptional profiling (**Suppl Table S1**). This then suggests that *MIR3607* indirectly promotes Pax6 translation, perhaps by blocking the post-transcriptional inhibition of Pax6 expression. In this line of evidence, our GSEA analyses revealed mTORC1 among the most significantly altered pathways upon *MIR3607* expression (NES: 2.22; *p*-val: 0.0078; *q*-val: 0.0054; **Suppl. Table S1**). Hyperactive mTORC1 is a shared molecular hallmark in several neurodevelopmental disorders characterized by abnormal brain cytoarchitecture. This is supported by evidence from multiple animal models suggesting a causative effect of increased mTORC1 signaling on abnormal cortical lamination, but much less is known about the underlying mechanisms (Feliciano et al., 2011; Kassai et al., 2014; Lin et al., 2016; Moon et al., 2015; Orlova et al., 2010; Tsai et al., 2014). Importantly, mTORC1 regulates Pax6 protein levels in the chick retina, where mTORC1 is activated downstream of Wnt/βCatenin signaling (Zelinka et al., 2016). Taken together, this suggests that in the developing cerebral cortex Pax6 protein levels are indirectly increased by *MIR3607* via augmenting Wnt signaling and then mTORC1 activity.

### *MIR3607* is a major regulator of the canonical Wnt/βCatenin signaling pathway

Expression of *MIR3607* in the early embryonic mouse cortex drives the amplification and massive delamination of apical progenitor cells to basal positions, frequently forming conspicuous proliferative rosettes. Our results also demonstrate that this complex phenotype is caused by the overactivation of the canonical Wnt/βCatenin pathway, and as such, it is largely rescued by expression of APC, which drives βCatenin for phosphorylation and proteasome degradation. Accordingly, the effects of expressing *MIR3607* in early mouse embryos were phenocopied by electroporation of constitutively active βCatenin, and are reminiscent of the alterations observed in transgenic mice expressing this construct in early cortical neuroepithelial cells (Chenn & Walsh, 2002; Chenn & Walsh, 2003). Our transcriptomic analyses demonstrate that *MIR3607* promotes canonical Wnt/βCatenin signaling at multiple levels, by targeting and significantly modifying mRNA levels of several major regulators of this pathway, including upregulation of *βCatenin* itself and downregulation of the pathway’s key repressor *APC*. Our study confirms previous reports showing a critical role of APC and Wnt signaling in progenitor cells during cerebral cortex development (Eom et al., 2014; Nakagawa et al., 2017; Yokota et al., 2009), and now demonstrates that *MIR3607* is a fundamental negative regulator of APC and activator of Wnt/βCatenin signaling in this process. This is in agreement with previous studies showing the regulatory interplay between *MIR3607* and APC to control the proliferative activity of the lung cancer cell line A549 (Lin et al., 2017).

A striking observation of our current study is that the effects of *MIR3607* on the embryonic development of the mouse cortex were very different if this was expressed starting at early (E12.5) or at mid (E14.5) stages of cortical development. The severe disruption of germinal layers caused when *MIR3607* was expressed starting at E12.5, remained ongoing by E15.5, and it eventually led to the formation of massive subventricular heterotopia in the postnatal mouse brain. In contrast, expression of *MIR3607* starting at E14.5 had a minor and very transient effect on progenitor cell amplification, which was surprising given the temporal overlap with the E12.5-E15.5 manipulations. Rather, the largest effect of late-onset *MIR3607* expression was on postmitotic neurons, altering their migration and laminar position as well as the growth of their axons towards the corpus callosum. This was consistent with part of the transcriptomic phenotype caused by *MIR3607* expression, with significant changes in gene networks related to mTORC1 signaling, axonal development and L1CAM interactions (Schmid & Maness, 2008).

While the origin of differences in phenotypes between early and mid developmental stages remains to be identified, existing evidence suggests that it may be related to a differential relevance of APC function along cortical development. It has been previously shown that the endogenous levels of βCatenin activity are downregulated in the embryonic cerebral cortex from E12.5 through E16.5 (Mutch et al., 2009). This suggests that Wnt/βCatenin activity may gradually become less relevant in progenitor cells as cortical development progresses. In contrast, expression of APC remains high at late stages (Yokota et al., 2009), which suggests that the endogenous downregulation of βCatenin activity is not related to increased APC expression. Hence, lowering APC levels by *MIR3607* may be less relevant at later stages, and thus have progressively less impact on progenitor cell amplification. Our results demonstrate the existence of epochs or temporal windows during cerebral cortex development with different sensitivity and responsiveness to specific genetic insults or manipulations, in our case the expression of *MIR3607*. We previously demonstrated a similar scenario in the ferret cerebral cortex, where dynamic changes in gene expression levels during development define a brief period of embryogenesis critical for aRGC delamination, genesis of bRGCs and formation of the OSVZ (Martinez-Martinez et al., 2016). Here, we further demonstrate that the severe disruption of cortical germinal layers at early stages caused by *MIR3607* expression and βCatenin overactivation follows a catastrophic dynamics, with these defects extending into late developmental stages including when the cortex would otherwise by largely insensitive to *MIR3607* expression. In summary, we have identified *MIR3607* as a major regulator of Wnt/βCatenin signaling, a fundamental pathway with key functions in the embryonic development of the cerebral cortex (Chenn & Walsh, 2002; Eom et al., 2014; Hirabayashi et al., 2004; Nakagawa et al., 2017; Yokota et al., 2009). Moreover, our study uncovers previously unrecognized critical roles for miRNAs in brain development (De Pietri Tonelli et al., 2008), regulating neural stem cell amplification and delamination, germinal layer stability, neurogenesis and axonal growth in the developing cerebral cortex. From the evolutionary point of view, our findings suggest that a loss of expression of *MIR3607* during cerebral cortex development may have been a key factor for the secondary reduction of brain size during recent rodent evolution.

## Materials and Methods

### Animals

Timed-pregnant sable ferrets (*Mustela putorius furo*) were obtained from Euroferret (Denmark) and maintained on a 16:8 hr light:dark cycle at the Animal Facilities of the Universidad Miguel Hernández. Wild-type mice in ICR background were obtained from a breeding colony at the Animal Facility of the Instituto de Neurociencias. The day a vaginal plug was detected was considered embryonic day (E) 0.5. Animals were treated according to Spanish (RD 53/2013) and European regulations, and experimental protocols were approved by the CSIC Ethics Committee.

### DNA constructs

For gain-of-function experiments, we used pCAG-GFP expressing green fluorescent protein under the CAG promoter, mixed with psil-pre-MIR3607 encoding *pre-MIR3607-5p* under the U6 promoter, or with psil-pre-miRScr under U6 as control. Oligos encoding pre-*MIR3607* (5’GATCCGACTGATTTCCTTCATGTCAAGCTTCAAGAGAGCATGTGATGAAGCAAATCAGTTTTTTT3’; SIGMA) were designed with BamHI-HindIII sites and cloned into pSilencer2.1-U6 puro (Thermo Fisher Scientific, Cat# AM5762). pSilencer puro negative control plasmid provided with the kit was used as psil-pre-miRScr, encoding an siRNA sequence not found in the mouse, human,or rat genome databases. Plasmid DNA was purified with a NucleoBond Xtra Midi kit (Cultech, 22740410.50) and resuspended in nuclease-free water (SIGMA). Other plasmids used were pCMV-GFP expressing GFP under the CMV promoter, pCMV-Neo-Bam APC (Addgene, #16507) and pCAG-Δ90βCateninGFP (Addgene, #26645) encoding stabilized βCatenin (Wrobel et al., 2007).

### Validation of *MIR3607* expression

Human embryonic kidney 293T (HEK293T) cells were transfected with 2µg of psil-pre-miRScr or psil-pre-MIR3607, plus 2µg of pCAG-GFP using Lipofectamine, harvested 2 days later, and RNA was isolated using *mir*Vana^TM^ miRNA isolation kit (Cat #AM1560). Quantitative RT-PCR was carried out using Taqman microRNA Assays. All kits and reagents, including primers and probes for *MIR3607* (Assay #463448) and U6 snoRNA control (Assay #001973) were from Thermo Fischer Scientific (Cat #4427975).

### *In utero* electroporation

*In utero* electroporation was performed as described elsewhere (Cardenas et al., 2018). Briefly, timed-pregnant females were deeply anesthetized with Isoflurane, the abdominal cavity was open and the uterine horns exposed. DNA solution (1μl) was injected into the lateral ventricle using pulled glass micropipettes, and square electric pulses (35V for E12.5, 45V for E14.5, 50ms on – 950ms off, 5 pulses) were applied with an electric stimulator (Cuy21EDIT Bex C., LTD) using round electrodes (CUY650P5, Nepa Gene). Plasmid concentrations for gain-of-function experiments were: GFP (0.7µg/µl), miRScr (1µg/µl), *MIR3607* (1µg/µl); for rescue experiments: miRScr (0.75µg/µl), *MIR3607* (0.75µg/µl), APC (0.75µg/µl), GFP (0.5µg/µl). Uterine horns were placed back into the abdominal cavity, suture closed, and the pregnant female was allowed to fully recover on a heating pad before returning to the home cage.

### hiPSC culture

Human iPSCs were cultured at 37°C, 5% CO_2_ and ambient oxygen level on Geltrex coated plates in mTeSR1 medium (STEMCELL Technologies, 05850) with daily medium change. For passaging, iPSC colonies were incubated with StemPro Accutase Cell Dissociation Reagent diluted 1:4 in PBS for 4 minutes. Pieces of colonies were washed off with DMEM/F12, centrifuged for 5min at 300 x g and resuspended in mTeSR1 supplemented with 10µM Rock inhibitor Y-27632(2HCl) for the first day.

### Generation of human cerebral organoids

Cerebral organoids were generated as previously described (Lancaster & Knoblich, 2014). Briefly, mycomplasma-free iPSCs were dissociated in to single cells using StemPro Accutase Cell Dissociation Reagent (A1110501, Life Technologies) and plated in the concentration of 9000 single iPSCs/well into low attachment 96-well tissue culture plates in hES medium (DMEM/F12GlutaMAX supplemented with 20% Knockout Serum Replacement, 3% ES grade FBS, 1% Non-essential amino acids, 0.1mM 2-mercaptoethanol, 4ng/ml bFGF and 50µM Rock inhibitor Y27632) for 6 days in order to form embryoid bodies (EBs). Rock inhibitor Y27632 and bFGF were removed on the 4th day. On day 6 EBs were transferred into low attachment 24-well plates in NIM medium (DMEM/F12GlutaMAX supplemented with 1:100 N2 supplement, 1% Non-essential amino acids and 5µg/ml Heparin) and cultured for additional 6 days. On day 12 EBs were embedded in Matrigel drops and then they were transfer in 10cm tissue culture plates in NDM minus A medium (DMEM/F12GlutaMAX and Neurobasal in ratio 1:1 supplemented with 1:100 N2 supplement 1:100 B27 without Vitamin A, 0.5% Non-essential amino acids, insulin 2.5µg/ml, 1:100 Antibiotic-Antimycotic and 50µM 2-mercaptoethanol) in order to form organoids. 4 days after Matrigel embedding cerebral organoids were transfer into an orbital shaker and cultured until electroporation in NDM plus A medium (DMEM/F12GlutaMAX and Neurobasal in ratio 1:1 supplemented with 1:100 N2 supplement 1:100 B27 with Vitamin A, 0.5% Non-essential amino acids, insulin 2.5µg/ml, 1:100 Antibiotic-Antimycotic and 50µM 2-mercaptoethanol). During the whole period of cerebral organoid generation, cells were kept at 37°C, 5% CO_2_ and ambient oxygen level with medium changes every other day. After transferring the cerebral organoids onto the shaker medium was changed twice per week.

### Electroporation of human cerebral organoids

Cerebral organoids were kept in antibiotics-free conditions prior to electroporation. Electroporations were performed in cerebral organoids at 37 days stages after the initial plating of the cells and fixed 7 days post electroporation. During the electroporation cerebral organoids were placed in an electroporation chamber (Harvard Apparatus, Holliston, MA, USA) under a stereoscope and using a glass microcapillary 1-2μl of plasmid DNAs was injected together with Fast Green (0.1%, Sigma) into different ventricles of the organoids. Plasmid DNA concentrations were: GFP (0.7µg/µl), miRScr (1µg/µl), *MIR3607* (1µg/µl). Cerebral organoids were subsequently electroporated with 5 pulses applied at 80V for 50ms each at intervals of 500ms (ECM830, Harvard Apparatus). Following electroporation, cerebral organoids were kept for additional 24hr in antibiotics-free media, and then changed into the normal media until fixation. Cerebral organoids were fixed using 4% PFA for 1hr at 4°C, cryopreserved with 30% sucrose and stored at −20°C. For immunofluorescence, 16μm cryosections were prepared.

### Bromodeoxyuridine labeling experiments

Bromodeoxyuridine (BrdU, SIGMA) solution (10mg/ml in 0.9% NaCl) was administered intraperitoneally at 50mg/kg body weight. To identify progenitor cells in S-phase, a single dose of BrdU was injected 30min before fixation. To measure cell cycle exit, a single dose of BrdU was administrated 24hr before fixation, and the percentage of GFP+/BrdU+/Ki67+ cells was measured.

### Tissue processing

Embryonic brains were immersion fixed in phosphate-buffered 4% paraformaldehyde (PFA) at 4°C. Postnatal pups were perfused transcardially with PFA and the heads were postfixed overnight at 4°C. After fixation, brains were washed with phosphate buffered saline (PBS) and cryoprotected overnight with 30% sucrose. For ISH, brains were treated under RNAse free conditions and cryoprotected in 30% sucrose, 2% PFA. Brains were finally cryosectioned at 20-30 µm.

### Immunohistochemistry

Brain sections were incubated with primary antibodies overnight, followed by appropriate fluorescently conjugated secondary antibodies and counterstained with 4′,6-diamidino-2-phenylindole (DAPI; Sigma, D9542). Primary antibodies used were against: BrdU (1:500, Abcam ab6326), GFP (1:1000, Aves Lab GFP-1020), Ki67 (1:300, Abcam ab15580), phosphohistone H3 (1:1000, Upstate 06-570), Tbr1 (1:500, Abcam ab31940), Tbr2 (1:500, Millipore ab31940), Par3 (1:500, Millipore MABF28), Pax6 (1:500, Millipore AB2237), Cux1 (1:500, Santa Cruz), activated β-Catenin (1:500, Merk 05-665). Secondary antibodies were from Vector Lab: biotinylated anti-Rat IgG (1:200, BA-9400), biotinylated anti-Rabbit IgG (1:200, BA-1000); from Jackson Inmunoresearch: Cy3 Fab fragment anti-Rabbit IgG (1:200, 711-167-003), Alexa488 anti-chicken IgY (1:200, 703-545-155), Cy5 Streptavidin (1:200, 016-170-084); from Invitrogen: Alexa555 anti-rabbit IgG (1:200, A-31572).

### miRNA *in situ* hybridization

miRNA *in situ* hybridization (ISH) was performed as described previously (De Pietri Tonelli et al., 2008). Cryostat, cryotome or deparafinized paraffin sections were washed briefly with PBS, permeabilized with RIPA buffer (150Mm Nacl, TritonX-100 1%, 0.5% sodium deoxycholate,0.1% SDS, 1 mM EDTA, 50Mm Tris), pre-hybridized at 54°C for 1 hr with hybridization solution (50% Formamide (Ambion), 5x SSC (Sigma), 5x Denhardt’s (from a 50x stock; Sigma), 250µg/ml of Yeast RNA, 500µg/Ml salmon sperm DNA) and hybridized overnight at 54°C (30°C below RNA Rm) with miRCURY™ LNA™ microRNA ISH Detection Probe (5’-3’-ACTGATTTGCTTCATCACATGC/3Dig_N/, product #612280-350 Exiqon) in hybridization solution. Sections were washed 2×1 hour with pre-warmed post-hybridization solution (50% Formamide, 2x SSC and 0.1% Tween), washed briefly with MABT buffer (1x maleic acid-Nacl pH 7.5, 0.1% Tween), blocked with blocking solution (10% Bovine serum in MABT buffer) for 1hr at room temperature, and incubated with alkaline phosphatase coupled anti-digoxigenin Fab fragments (1:2000, Roche) in blocking solution overnight at 4°C. After brief washes with MABT and NTMT (100Mm Tris pH 9.5, 50Mm MgCl_2,_ 100Mm NaCl, 0.1% Tween), sections were incubated in nitroblue tetrazolium (NBT)/ 5-bromo-4-chloro-3-indolyl phosphate (BCIP) solution [3.4 µg/ml from NBT stock and 3.5 µl/ml from BCIP stock in NTMT buffer (100 mg/ml NBT stock in 70% dimethylformamide, 50mg/ml BCIP stock in 100% dimethylformamide; Roche)]. Sections were finally dehydrated and coversliped.

### FACS sorting of electroporated brains

Electroporated mouse embryos with similar rostro-caudal and latero-medial location of GFP fluorescence in the neocortex were selected and the GFP+ cortical area was dissected out in ice-cold HBSS (Thermo Fisher Scientific). Dissected tissue was dissociated with Trypsin diluted in HBSS in the presence of DNAse at 37°C for 8 minutes. HBSS supplemented with 10% FBS and 1% penicillin/streptomycin (P/S) was then added, and tissue was gently triturated. The cell suspension was centrifuged, and cells in pellet were resuspended in 500µl of medium, filtered and FACS sorted (FACS Aria II, BD). Cells with high GFP intensity were collected in Neurobasal medium supplemented with 10% FBS, and 1% P/S, and RNA was extracted using Arcturus PicoPure^TM^ RNA isolation kit (Thermo Fischer Scientific, Cat # KIT0202) according to the manufactureŕs protocol. RNA integrity was analysed using Bioanalyzer (Agilent 2100) and samples with RIN values above 9.5 were selected for RNA sequencing. For each biological replica, cells from two embryos were pooled together, and three independent replicas per condition were sequenced. Libraries were prepared with the SMART-seq v4 Library Prep Kit and sequenced on Illumina HISeq 2500 sequencer using 50bp single reads.

### RNA sequencing and differential expression analysis

Sequencing reads were aligned using HISAT2 v2.1.0 (Kim et al., 2015) to GRCm.38/mm10 mouse genome. Integrative Genomics Viewer (Robinson et al., 2011) was used to visualize aligned reads and normalized coverage tracks (RPM). Reads overlapping annotated genes (Ensemble GRCm38.93) were counted using HTSeq v0.11.1 (Anders et al., 2015). Differentially expressed genes (DEGs) were detected using DESeq2 v1.18.1 package (Love et al., 2014) in R (Ihaka & Gentleman, 1996). Genes with Adj. *p* < 0.01 were considered significantly differentially expressed. For some comparisons, mRNA abundance of RNA-seq data was obtained using transcript per million normalization (TPM) (Wagner et al., 2012). RNA-seq data reported in this study are accessible through the Gene Expression Omnibus (GEO) database with the GEO Series accession number GSE135321.

### Gene set enrichment and pathway analyses

Identification of enriched biological functions and processes in DEGs was performed using clusterProfiler (Yu et al., 2012) package in R. Over-representation test (Boyle et al., 2004) was performed using enrichGO function of clusterProfiler R package with the following parameters: gene=DEGs; OrgDb=org.Mm.eg.db; ont=BP; pAdjustMethod=BH; pvalueCutoff=0.05; qvalueCutoff=0.1). Functional Annotation Analysis of DEGs was performed using DAVID v6.8 (Huang da et al., 2009) with the following annotations: 3 Functional Categories (COG_ONTOLOGY, UP_KEYWORDS, UP_SEQ_FEATURE); 3 Gene Ontologies (GOTERM_BP_DIRECT, GOTERM_CC_DIRECT, GOTERM_MF_DIRECT); 2 Pathways (BIOCARTA, KEGG_PATHWAY); 3 Protein Domains (INTERPRO, PIR_SUPERFAMILY, SMART). Visualization of interrelations of terms and functional groups in biological netwoks was perfomed using ClueGO v2.5.0 (Bindea et al., 2009) plug-in of Cytoscape v3.6.0 (Shannon et al., 2003) with the following parameters: gene list=DEGs; ontologies= GO_MolecularFunction-EBI-QuickGO-GOA_22.03.2018_00h00, GO_BiologicalProcess-EBI-QuickGO-GOA_22.03.2018_00h00, WikiPathways_10.01.2019, KEGG_10.01.2019, REACTOME_Reactions_10.01.2019, REACTOME_Pathways_10.01.2019; Statistical Test Used = Enrichment/Depletion (Two-sided hypergeometric test); Correction Method Used = Bonferroni step down; Min GO Level = 3; Max GO Level = 8; Number of Genes = 3; Min Percentage = 4.0; Combine Clusters With ‘Or’ = true; Percentage for a Cluster to be Significant = 60.0; GO Fusion = true; GO Group = true; Kappa Score Threshold = 0.4; Over View Term = SmallestPValue; Group By Kappa Statistics = true; Initial Group Size = 1; Sharing Group Percentage = 50.0. Gene Set Enrichment Analysis (GSEA) (Subramanian et al., 2005) was performed using GSEAPreranked tool and GSEA function on clusterProfiler package in R, on 21822 unique features (genes) that were pre-ranked on the log2 of the fold-change from DESeq2 analysis. The following parameters were used for GSEA: exponent = 0 (scoring_scheme=classic), nPerm=1000, minGSSize=15, maxGSSize=500. GSEA was performed using MSigDB gene sets Hallmark (h.all.v6.2.symbols.gmt) and GO (c5.all.v6.2.symbols.gmt).

### *MIR3607* target prediction

miRNA targets were predicted using available online tools: TargetScanHuman 7.2, miRDB (Wong & Wang, 2015), TargetScanMouse Custom (Version 4 and 5.2) and miRDB custom prediction for identification of mouse target genes.

#### Image analysis, quantification and statistics

Images were acquired using a florescence microscope (Zeiss Axio Imager Z2) with Apotome.2 and coupled to two different digital cameras (AxioCam MRm and AxioCam ICc) or an inverted confocal microscope (Olympus FluoView FV1000). All images were analysed with ImageJ (Fiji). Co-localization studies were performed on single plane confocal images from 40X Z stacks. Cell distribution analyses were performed using Neurolucida and Neuroexplorer software (MBF Bioscience). All quantifications were performed at the same rostral-caudal and latero-medial levels on at least three different embryos from at least two independent litters. ISH images were acquired using Zeiss Axio Imager Z2. Brightness and contrast of the images shown in figures were homogeneously adjusted for clarity using Adobe Photoshop. Statistical analyses were carried out in Microsoft Excel or GraphPad Software using ANOVA with post-hoc Bonferroni correction (equal variances) or the Welch test with post-hoc Games-Howell (different variances), Kolmogorov-Smirnov test, χ^2^-test or independent samples *t*-test, where appropriate and upon normality testing. Significance was set at *p*=0.05. In the analyses of Pax6 and Tbr2 co-expression, the influence of each protein’s increased abundance over their increased co-expression in *MIR3607*-expressing embryos was tested by mathematical correction. This was performed by dividing the Pax6 intensity value of each cell by the average increase in Pax6 intensity between *MIR3607* and *Scr* embryos. The same method was used for Tbr2.

## Acknowledgements

We thank G. Exposito, V. Villar and A. Caler for excellent assistance with imaging and FACS, A. Espinós and J. Brotons-Mas for help with statistics, and M. Drukker for hiPSCs. We also thank members of our lab for insightful discussions and critical reading of the manuscript. K.C. was recipient of a Santiago Grisolía fellowship, A.P.-C. was recipient of a predoctoral fellowship from Fundación Tatiana Pérez de Guzmán el Bueno. This work was supported by grants to V.B. from the Spanish State Research Agency (SAF2015-69168-R, PGC2018-102172-B-I00) and European Research Council (309633). J.L-A. was supported by grants RYC-2015-18056 and RTI2018-102260-B-100 from Spanish State Research Agency co-financed by ERDF. V.B. acknowledges financial support from the Spanish State Research Agency, through the “Severo Ochoa” Programme for Centers of Excellence in R&D (ref. SEV-2017-0723).

## Author contributions

K.C. and V.B. conceived and designed the experiments; K.C., A.P.-C., Y.N. and A.C. performed experiments; K.C., A.M.-G., A.P.-C., A.C., Y.N. and V.B. analyzed experiments; J-P.L.A. and V.B. provided reagents and resources, supervised experiments and analyses. V.B. provided funding and wrote the manuscript with input from the other authors.

